# Rbm8a deficiency causes hematopoietic defects by modulating Wnt/PCP signaling

**DOI:** 10.1101/2023.04.12.536513

**Authors:** Agnese Kocere, Elena Chiavacci, Charlotte Soneson, Harrison H. Wells, Kevin Manuel Méndez-Acevedo, Jacalyn S. MacGowan, Seth T. Jacobson, Max S. Hiltabidle, Azhwar Raghunath, Jordan A. Shavit, Daniela Panáková, Margot L. K. Williams, Mark D. Robinson, Christian Mosimann, Alexa Burger

## Abstract

Defects in blood development frequently occur among syndromic congenital anomalies. Thrombocytopenia-Absent Radius (TAR) syndrome is a rare congenital condition with reduced platelets (hypomegakaryocytic thrombocytopenia) and forelimb anomalies, concurrent with more variable heart and kidney defects. TAR syndrome associates with hypomorphic gene function for *RBM8A/Y14* that encodes a component of the exon junction complex involved in mRNA splicing, transport, and nonsense-mediated decay. How perturbing a general mRNA-processing factor causes the selective TAR Syndrome phenotypes remains unknown. Here, we connect zebrafish *rbm8a* perturbation to early hematopoietic defects via attenuated non-canonical Wnt/Planar Cell Polarity (PCP) signaling that controls developmental cell re-arrangements. In hypomorphic *rbm8a* zebrafish, we observe a significant reduction of *cd41*-positive thrombocytes. *rbm8a*-mutant zebrafish embryos accumulate mRNAs with individual retained introns, a hallmark of defective nonsense-mediated decay; affected mRNAs include transcripts for non-canonical Wnt/PCP pathway components. We establish that *rbm8a*-mutant embryos show convergent extension defects and that reduced *rbm8a* function interacts with perturbations in non-canonical Wnt/PCP pathway genes w*nt5b*, *wnt11f2*, *fzd7a*, and *vangl2*. Using live-imaging, we found reduced *rbm8a* function impairs the architecture of the lateral plate mesoderm (LPM) that forms hematopoietic, cardiovascular, kidney, and forelimb skeleton progenitors as affected in TAR Syndrome. Both mutants for *rbm8a* and for the PCP gene *vangl2* feature impaired expression of early hematopoietic/endothelial genes including *runx1* and the megakaryocyte regulator *gfi1aa*. Together, our data propose aberrant LPM patterning and hematopoietic defects as consequence of attenuated non-canonical Wnt/PCP signaling upon reduced *rbm8a* function. These results also link TAR Syndrome to a potential LPM origin and a developmental mechanism.

**HIGHLIGHTS:**

- Zebrafish mutants for the TAR Syndrome gene *rbm8a* show thrombocyte reduction
- Attenuated Rbm8a function results in retained introns in mRNAs encoding PCP components
- Early PCP defects result in lateral plate mesoderm anomalies and hematopoietic defects
- PCP anomalies selectively impact cell fate patterning

## INTRODUCTION

Mutations in genes that lead to pleiotropic phenotypes, as observed in complex syndromic birth anomalies, remain challenging to mechanistically connect to a tissue- or cell type-specific developmental defect. Thrombocytopenia with absent radius (TAR) Syndrome (OMIM #274000) is a rare (0.42 per 100’000 live births), autosomal-recessive congenital disorder that at birth manifests as blood platelet deficiency (hypomegakaryocytic thrombocytopenia) and bilateral absence of the radius bone ^1,2^. While thrombocytopenia is the most consistent phenotype, the limb defects vary considerably among patients and can involve both arms and legs. Concomitant anomalies in TAR syndrome patients can include heart malformations, such as Tetralogy of Fallot or atrial septal defects, floating or unilaterally absent kidneys, as well as mild craniofacial alterations ^2–5^. Curiously, the predominant coagulation defect of newborn TAR patients commonly recedes with age, and patients show normal platelet counts and blood clotting responses a few years after birth ^1,2^. The complex combination of organs and cell types affected in TAR patients has so far precluded any clear assignment of a developmental cause underlying the syndrome.

Genetically, *RBM8A* has been identified as causative gene in TAR syndrome patients. Sequence analysis of TAR patients has revealed that compound inheritance of a *1q21.1* deletion that includes *RBM8A* is required with a noncoding single-nucleotide variant in the remaining copy of the *RBM8A* gene ^6^. Corroborating the causal link between TAR Syndrome and *RBM8A*. The major *RBM8A* allele combinations in TAR patients suggest that one gene copy is a complete null by deletion and the other is hypomorphic, while individual patients with bi-allelic *RBM8A* variants in the absence of *1q21.1* deletions have been reported ^1,2,5–7^. *RBM8A* encodes RBM8A/Y14 that together with eIF4A-III, MLN51, and MAGOH constitutes the exon–junction complex (EJC) ^8^. The EJC is involved in essential post-transcriptional mRNA control by associating with exon-exon junctions after splicing to support nuclear export and sub-cellular localization of specific transcripts, translation enhancement, and nonsense-mediated RNA decay (NMD) ^9^. NMD as major mRNA quality control step during translation leads to degradation of mRNAs with retained introns as routinely caused by incomplete splicing, in particular of long introns ^10–13^. Which mechanism(s), mRNA target(s), and developmental process(es) are affected in TAR Syndrome upon reduced RBM8A function remain unknown.

Pioneering work on recessive zebrafish *rbm8a* and *magoh* mutants has documented the impact of EJC perturbation in which maternal mRNA and protein contribution to the embryo rescues development until lack of zygotic supply causes aberrant phenotypes ^14^. Zygotic *rbm8a* loss-of-function resulted in no discernible morphological phenotypes until late segmentation stages (approximately 19 hpf), when cell death in the brain and general muscle paralysis became detectable ^14^. Notably, zebrafish *rbm8a* mutants showed disrupted NMD and increased stability of mRNAs with 3’ UTR introns by 24 hpf ^14^. In *Drosophila*, tissue-specific disruption of NMD using *rbm8a* and *magoh* mutants has been linked to accumulation of individual mRNAs with retained large introns that result in perturbation of select developmental processes including RAS/MAPK signaling ^10,11^. Together, these data document both quantitative as well as qualitative impact on individual mRNAs upon EJC perturbation in different models.

A possible lead to connect the seemingly pleiotropic TAR Syndrome phenotypes comes from the developmental origin of the affected organs. The lateral plate mesoderm (LPM) forms the progenitor cells for blood, heart, vasculature, kidney, craniofacial muscles, and limb connective tissue in the developing vertebrate embryo ^15–20^. Hypomorphic perturbation of *RBM8A* in TAR Syndrome could therefore impact genes involved in a shared developmental process at the base of blood, limb, heart, and kidney development. The diversity of LPM cell fates could explain complex, yet developmentally connected co-morbidities of syndromic congenital anomalies as LPM diseases that result from early patterning defects^21^. By affecting blood, forelimbs, heart, and kidneys, the spectrum of TAR syndrome phenotypes seems to predominantly affect tissues of LPM origin. Yet, what developmental mechanisms altered RBM8A/Y14 dosage in TAR Syndrome triggers to possibly cause a LPM defect, and in particular thrombocytopenia, remains unknown.

Hematopoietic cell lineages emerge in close association with endothelial progenitors within the LPM from bilateral progenitors expressing the transcription factors Tal1/Scl, Lmo2, and Etv2, starting with a first wave of *Gata1*-expressing primitive erythrocytes ^19,22–25^. Subsequently emerging intermediate hematopoietic progenitors form primitive myeloid cells including transient megakaryocytes that in mammals begin to shed anucleate thrombocytes (also called platelets) for coagulation ^26–29^. The *Runx1*-expressing hematopoietic stem cell (HSC) precursors emerge from the ventral wall of the dorsal aorta and undergo various maturation steps before seeding the final hematopoietic niches ^30–33^. Definitive megakaryocytes emerge from HSC-derived common myeloid progenitors that also form erythrocytes plus mast cells and myeloblasts ^25,34–36^. The Gfi1 transcription factors have been implicated in controlling hematopoietic lineage progression in human and mouse: Gfi1b predominantly drives megakaryocyte differentiation by suppressing erythroid fate, with missense mutations in human *GFI1B* causing thrombocytopenia ^37–39^. In zebrafish, the orthologs *gfi1aa* and *gfi1b* contribute to primitive erythropoiesis and are expressed in erythroid, intermediate erythroid-myeloid progenitors (EMPs), and emerging hematopoietic stem cells ^40–42^. At which step embryonic perturbation of Rbm8a affects thrombocyte formation and how it connects to the other LPM-associated defects awaits clarification.

Among the highly dynamic LPM that patterns while the embryo undergoes dramatic changes in length and cell number, hematopoietic progenitors emerge under the influence of general patterning and morphogenetic signals. Most-prominently, BMP and canonical Wnt/beta-catenin signaling have been linked to various steps in LPM patterning and hematopoietic lineage differentiation ^20,25,43–47^. In contrast, non-canonical Wnt/Planar cell polarity (PCP) signaling is a critical pathway coordinating the polarized orientation of fields of cells and relative orientation of cells among their neighbors during early embryo morphogenesis ^48–50^. Wnt/PCP signaling coordinates the convergent extension of the embryo during somite stages that also influence the lateral-to-medial migration of the LPM ^51–54^. Triggered by ligands including Wnt5 and Wnt11, select Frizzled receptors together with Celsr/Flamingo co-receptors and Vangl1/2 relay ligand binding through cytoplasmic components including Prickle1 and Dishevelled to control cytoskeletal dynamics without apparent transcriptional response ^49^. Beyond gross trunk defects resulting from global perturbation of the pathway, more selective defects in non-canonical Wnt/PCP signaling have been linked to structural anomalies affecting the neural tube, kidneys, and heart ^51,52,54–61^. Non-canonical Wnt signaling has been implicated in supporting hematopoietic stem cell emergence from the dorsal aorta by activating Notch ligands in somites ^62,63^, and in the maintenance of hematopoietic stem cells in their bone marrow niche ^64^. How deregulation of non-canonical Wnt/PCP signaling could contribute to earliest blood progenitor formation, migration, and congenital hematopoietic disease remains unknown.

Here, harnessing the dosage range provided by maternal contribution, different mutant *rbm8a* alleles, and antisense morpholino knockdown in zebrafish, we investigate if Rbm8a perturbation akin to TAR Syndrome causes the thrombocytopenia phenotype by impacting hematopoietic development from LPM. We provide evidence that reduced *rbm8a* function causes intron retention and misexpression of mRNAs encoding components of the non-canonical Wnt/PCP pathway. Subsequently, *rbm8a* attenuation results in several, cumulatively significant defects in early LPM and endothelial/hematopoietic lineage patterning that we also document in embryos with classic PCP defects. Our data connect impaired PCP signaling with hematopoietic phenotypes as possible developmental origin of the phenotypes observed in TAR Syndrome.

## RESULTS

### Null and hypomorphic *rbm8a* perturbation in zebrafish

Pioneering previous work has established that zebrafish *rbm8a* is maternally contributed as mRNA and protein, persisting for at least 24 hpf ^14^. This waning maternal contribution has been used to assess hypomorphic and loss-of-function phenotypes from 19 hours post fertilization (hpf) on, yet *rbm8a*-mutant embryos deteriorate rapidly past 24-28 hpf and are severely deformed by 72 hpf ^14^. We therefore sought genetic means to decrease Rbm8a protein levels to establish hypomorphic Rbm8a function beyond these timepoints.

To generate mutant alleles, we targeted the zebrafish *rbm8a* locus by CRISPR-Cas9 mutagenesis to induce non-homologous end joining (NHEJ)-mediated lesions in the coding sequence (**Fig. 1A**). We raised surviving low-dose crispants and isolated two different mutant *rbm8a* alleles: *g.5152_5156del* and *rbm8a g.5152_5154del*, subsequently abbreviated as *rbm8a^Δ5^* and *rbm8a^Δ3^*, respectively (**Fig. 1A**). *rbm8a^Δ5^* is a frameshift allele with no downstream alternative start codons that likely represents a null allele (**Fig. 1A**); no Rbm8a protein is detected by Western Blot at 24-26 hpf after the considerable maternal mRNA and protein deposition subsides (**Fig. 1B, Supplementary Fig. 1A-D**). Notably, our independently isolated allele is molecularly identical to the allele previously reported by Gangras and colleagues and generated with the same recommended sgRNA sequence, indicating preferential repair of this lesion with a 5 bp deletion ^14^. In contrast, our additional allele *rbm8a^Δ3^* deleted three base pairs, replacing *phenylalanine* (*F*) and *proline* (*P*) codons with a single *serine* (*S*) codon, substituting two non-polar side chains with a polar OH group (**Fig. 1A**). *rbm8a^Δ3^* substitutes amino acids right before the first alpha-helix in the Rbm8a N-terminus that is in proximity with the beta-sheet in Magoh ^65–67^, proposing *rbm8a^Δ3^* as potential hypomorphic allele.

**Figure 1:**
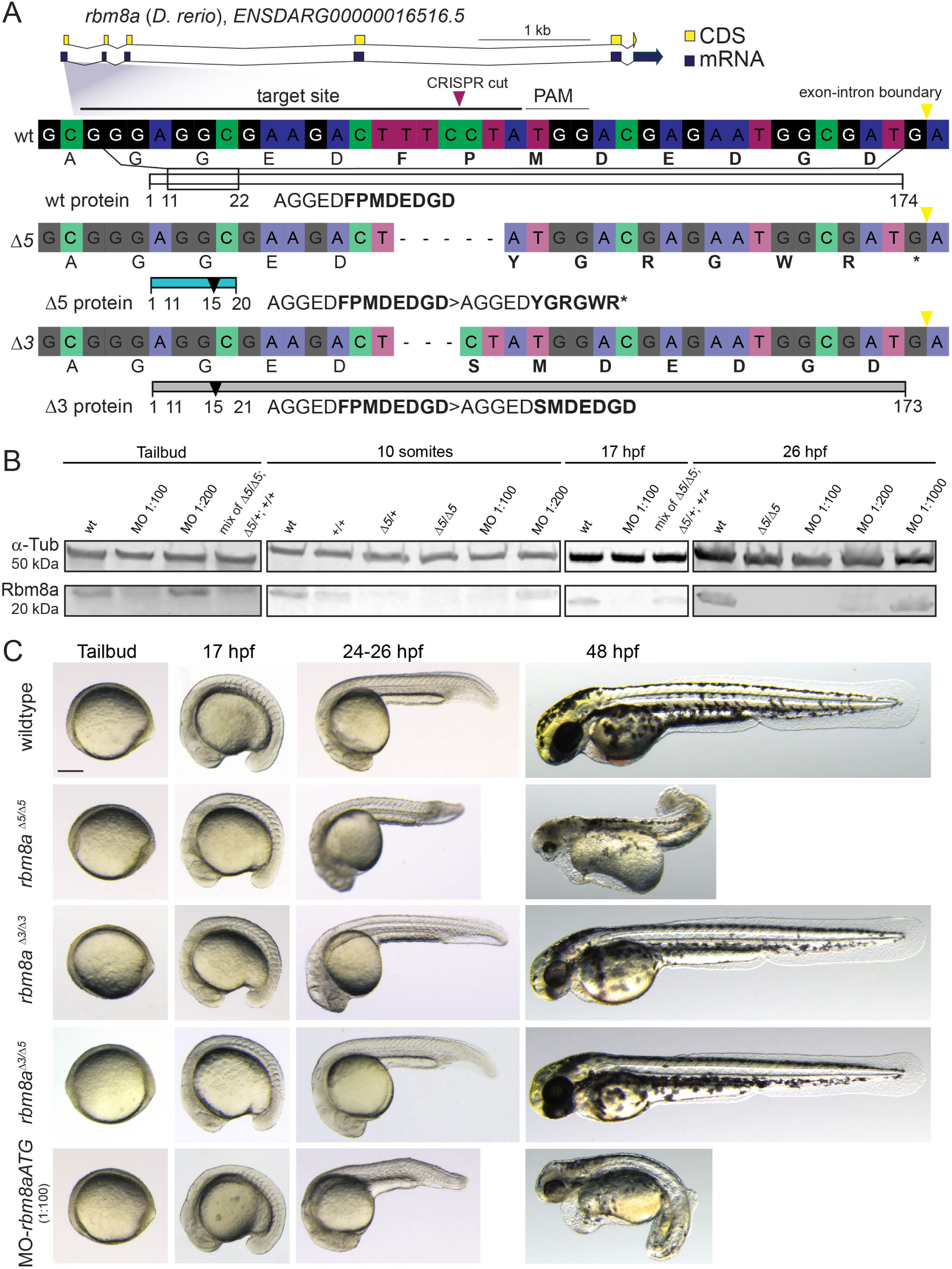
Genetic perturbations in the zebrafish *rbm8a* gene. (**A**) Allelic series of *rbm8a* in zebrafish with wild type allele, *rbm8a^Δ5^* allele and *rbm8a^Δ3^* allele induced by Cas9-mediated mutagenesis. (**B**) Western blot analysis of Rbm8a protein in mutants and morphants at different developmental stages, showing gradual decrease of maternal protein in mutants and translation block in morphants. (**C**) Representative images of zebrafish embryos carrying different *rbm8a* allele combinations during early development. Compared to wild type and different *rbm8a^Δ5^* and *rbm8a^Δ3^* combinations, homozygous *rbm8a^Δ5/Δ5^* carriers display severe microcephaly and corkscrew tail phenotype. *rbm8a* morphant (1:100 morpholino dilution) phenotypically recapitulates the homozygous *rbm8a^Δ5/Δ5^* embryos. Scale bars in (**C**): 250 μm, applies to all panels in C.

Embryos heterozygous for either allele (*rbm8a^Δ3/+^* and *rbm8a ^Δ5/+^*) showed no obvious malformations at observed timepoints up to 5 days post-fertilization and adults were viable and fertile. As previously reported and compared to wild type siblings, homozygous *rbm8a^Δ5^* (*rbm8a^Δ5/Δ5^*) zebrafish developed seemingly normal for the first 19 hpf, with subtle axis shortening during segmentation, before rapidly developing severe body axis hypoplasia (**Fig. 1C**). Around 26 hpf, the embryos started to display hypoplastic heads and tails with twisted axis, yet featured slowed heartbeat and rudimentary circulation before progressively deteriorating without survival past 60-72 hpf (**Fig. 1C**). In contrast, homozygous *rbm8a^Δ3^* (*rbm8a^Δ3/Δ3^*) embryos developed and survived without obvious overt defects into fertile adults (**Fig. 1C**). By crossing, we derived trans-heterozygous *rbm8a^Δ5/Δ3^* embryos that harbor one allele as likely null while the other provides hypomorphic activity; *rbm8a^Δ5/Δ3^* embryos resembled wild type siblings during early development and continued to develop seemingly normal until adulthood. Incrosses of adult *rbm8a^Δ5/Δ3^* resulted in the expected phenotypes and allele combinations, as observed in heterozygous incrosses of the individual alleles.

To further control embryonic Rbm8a protein levels, we obtained the translation-blocking morpholino *MO-rbm8aATG*. Injection of 1 nL of 0.01 nmol/µL *MO-rbm8aATG* phenocopied *rbm8a^Δ5/Δ5^* embryos by morphology (**Fig. 1C**). Morpholino injection also resulted in rapid translation block and reduction of Rbm8a protein already at tailbud stage and absence by 17 hpf, as detected by Western blot (**Fig. 1B, Supplementary Fig. 1A-D**). Co-injection of *MO-rbm8aATG* with capped mRNA encoding human *RBM8A* that has no sequence overlap with the morpholino rescued the knockdown embryos to at least 5 dpf, while *EGFP* mRNA as control had no rescue capacity (**Supplementary Fig. 1E-K, Supplementary Data 1**). In sum, these data validate *MO-rbm8aATG* as complementary tool to our *rbm8a* allelic series^68^. Altogether, these reagents establish a zygotic-mutant allelic series of zebrafish *rbm8a* and a potent translation-blocking knockdown reagent to study the impact of *rbm8a* perturbations on early development (**Supplementary Fig. 1L**).

To gain insights into specific phenotypes in our mutants, we used mRNA *in situ* hybridization (ISH) for diverse markers related to TAR Syndrome-associated structural defects (**Supplementary Fig. 2A**). We did not observe any overt phenotypes in *rbm8a*-mutant embryos at 17 hpf (around 14-15 somite stage), 48 hpf, and 72 hpf (**Supplementary Fig. 2B-D**): we observed no striking differences in the expression patterns of cardiovascular and hematopoietic (*pu.1, fli1a*, *vcana*, *gata1*), mesothelial (*hand2, wt1a*), kidney (*pax2a, wt1a*) and broader LPM (*bmp4*) markers in *rbm8a^Δ5/Δ5^* or *rbm8a^Δ3/Δ5^* embryos compared to wild type at 14-15 somite stage, despite the increasing deterioration in overall morphology observed in *rbm8a^Δ5/Δ5^* ^14^. We noted in individual *rbm8a^Δ5/Δ5^* embryos at 17 hpf that expression of the LPM genes *hand2* and *gata1* showed short, subtle interruptions in the continuous bilateral expression domains, which were however of variable penetrance among embryos (**Supplementary Fig. 2B**; see also below). Based on this first, albeit limited, gene expression analysis, we conclude that *rbm8a*-mutant zebrafish do not feature overt phenotypes in several TAR Syndrome-associated tissues or cell types at early stages of development, including the forelimbs (pectoral fins), kidney, or circulatory system. These observations are consistent with residual maternal contribution of Rbm8a-encoding mRNA and protein in our zygotic mutant conditions (**Fig. 1**)^14^.

### Hypomorphic *rbm8a* expression results in thrombocyte reduction

The functional equivalent of cytoplasmic platelets shed by megakaryocytes in humans, nucleated thrombocytes in zebrafish are labeled by the transgenic reporter *cd41:EGFP* starting from 40-46 hpf^69,70^. Notably, *cd41:EGFP* transgenics harbor EGFP-high thrombocytes and EGFP-low prospective hematopoietic stem cell precursors (HSPCs) that can be distinguished by sorting and fluorescence imaging^71,72^. As *rbm8a ^Δ5/Δ5^* mutants and *MO-rbm8aATG* morphants deteriorate beyond 28-30 hpf (**Fig. 1C**) ^14^, we sought to test thrombocyte numbers in genetic combinations with the hypomorphic *rbm8a^Δ3^* allele. We additionally established an injection titration curve for *MO-rbm8aATG* to reduce Rbm8a protein levels and to maintain viability: the suboptimal dose ranged between 1:125 and 1:200 stock dilution and resulted in slight to no detectable developmental defects in the morphants with survival beyond 7 dpf, compared to a dose of 1:100 that fully phenocopied *rbm8a^Δ5/Δ5^* mutants (**Fig. 1C**).

We next sought to quantify thrombocytes in wild type controls and upon mutant and morpholino-induced *rbm8a* perturbation in *cd41:EGFP*-transgenic larvae at 3 dpf and 6 dpf, when thrombocytes are clearly detectable. Previous work had applied live-imaging of circulating *cd41:EGFP* cells on a chosen field of view and counted cells during a pre-determined time window in the resulting video capture^73,74^. To increase throughput and to capture all *cd41:EGFP*-positive thrombocytes in individual larvae, we devised an alternative imaging-based workflow. To transiently arrest the heartbeat and stop circulation, we treated the larvae with 2,3-Butanedione monoxime (BDM) and quantified EGFP-positive cells from fluorescent images of whole larvae using custom Fiji scripts (**Fig. 2A**). This approach provided a semi high-throughput quantification to distinguish the circulating, high EGFP-expressing thrombocytes from the immobile, sparser low EGFP-expressing HSPCs in the caudal hematopoietic territory (CHT) (**Fig. 2A-I**). Our quantification documented a significant decrease in the number of high EGFP-positive cells at both 3 and 6 dpf in the morphants with a clear response to the dose of morpholino compared to wild type animals (**Fig. 2B-E,J,K, Supplementary Data 2**). Notably, viable mutant allele combinations for *rbm8a* showed no significant change to *cd41:EGFP*-expressing cell numbers at 3 dpf (**Fig. 2F,H,J, Supplementary Data 2**), in line with unperturbed maternal deposition of *rbm8a* mRNA and translated protein (**Fig. 1C**) ^14^. In contrast, at 6 dpf, *rbm8a^Δ3^*-homozygous larvae showed a significant reduction of *cd41:EGFP* cell counts, while *rbm8a^Δ3/Δ5^* larvae showed an even further reduction (**Fig. 2G,I,K, Supplementary Data 2**). This reduction was significant in larvae derived from incrosses of *rbm8a^Δ3/Δ5^* parents in which the oocytes only harbor the hypomorphic *rbm8a* allele as functional copy, further underlining the influence of maternally contributed wild type *rbm8a* transcript. These data are in line with a potential hypomorphic quality of the *rbm8a^Δ3^* allele. Of note, counting thrombocytes at 6 dpf using video-based methodology as per previous work ^73^ resulted in comparable observations and trends: wild type and heterozygous *rbm8a^Δ5/+^* larvae showed comparable numbers to the semi-automated count (**Supplementary Fig. 2E, Supplementary Data 2**). Homozygous *rbm8a^Δ3/Δ3^* and trans-heterozygous *rbm8a^Δ3/Δ5^* larvae showed slightly lower overall *cd41:EGFP*-expressing cell numbers using the video-based count, which further resulted in mild, but not significant, reduced thrombocyte numbers in the homozygous *rbm8a^Δ3/Δ3^* and trans-heterozygous *rbm8a^Δ3/Δ5^* larvae compared to wild type (**Supplementary Fig. 2F, Supplementary Data 2**). Together, our data indicate that reducing *rbm8a* function in zebrafish causes a reduction in *cd41:EGFP*-expressing thrombocytes.

**Figure 2:**
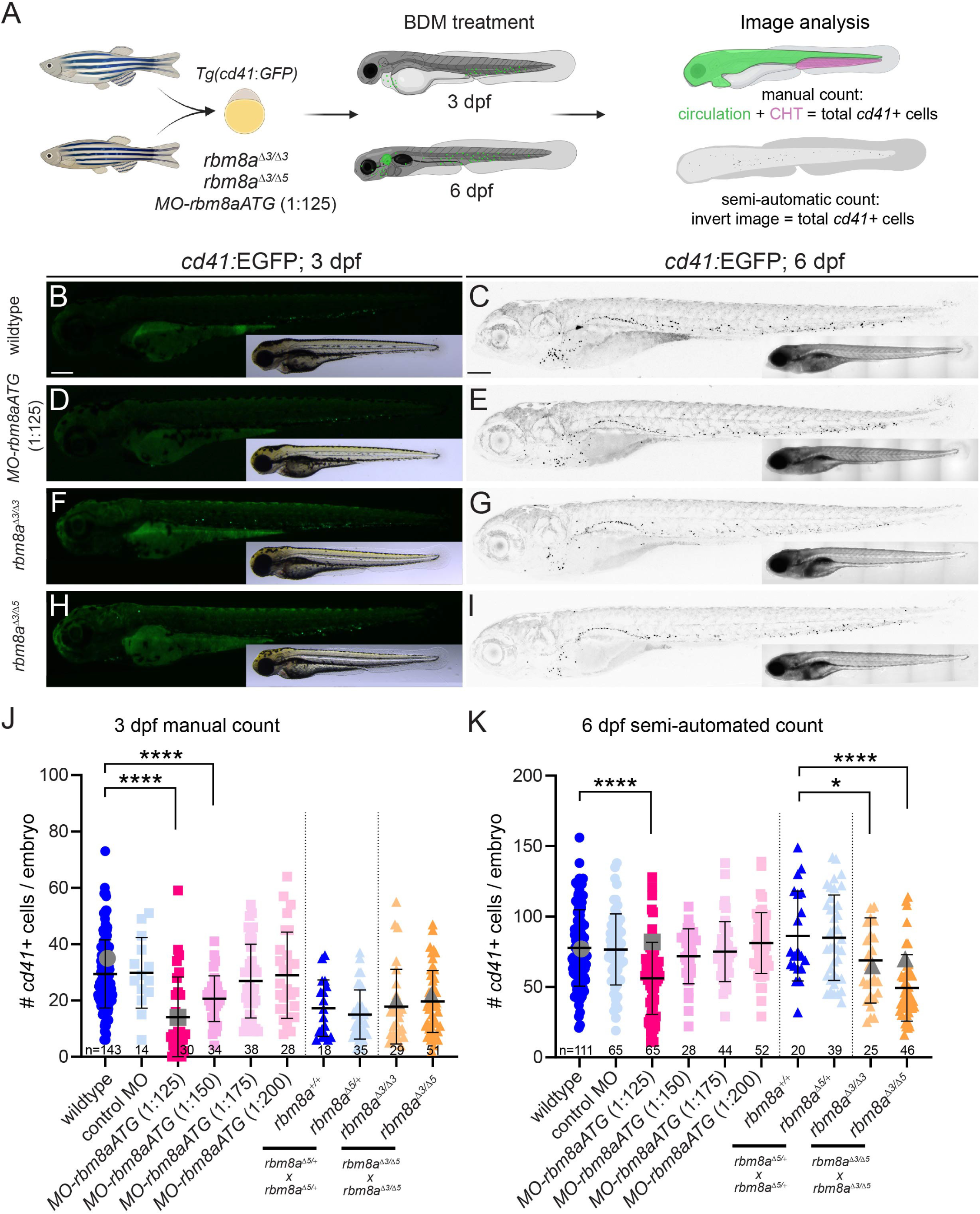
*rbm8a* perturbation reduces thrombocyte numbers in zebrafish larvae. (**A**) Workflow schematic of *cd41:*EGFP-positive thrombocyte progenitor quantification from circulation and from the caudal hematopoietic territory (CHT). Created with Biorender.com (Subscription: Individual, Agreement number: KV25WQCJKF). (**B**-**I**) Representative fluorescent dissecting scope and confocal images of zebrafish embryos transgenic for the thrombocyte marker *cd41:EGFP*, greyscale and color-inverted to reveal GFP-positive cells; anterior to the left, insert depicts brightfield image of larvae for reference. (**J**,**K**) *cd41:*EGFP-positive thrombocyte counts at 3 dpf (**J**) and 6 dpf (**K**) for each analyzed condition. Wild type larvae have significantly more *cd41*:EGFP-positive cells at 3 and 6 days compared to the high morpholino dose (1:125 and 1:150, only at 3 dpf), while lower dose (1:200) remained at wild type levels (**J**,**K**). At 6 dpf (**K**), viable trans-heterozygous allele combinations for *rbm8a* also show a significant thrombocyte reduction. Note that morpholino injections will block translation from maternal mRNA, while mutant combinations will retain maternal mRNA function from the wild type allele carried by the mother. Individual datapoints (total number of *cd41:GFP*-positive cells per embryo) shown with mean and standard deviation, significance calculated by Mann-Whitney test: 3 dpf (**J**) wild type vs. control MO p=0.7756 (not significant), wild type vs. *MO-rbm8aATG* 1:125 p<0.0001, wild type vs. *MO-rbm8aATG* 1:150 p<0.0001, wild type vs. *MO-rbm8aATG* 1:175 p=0.2589 (not significant), wild type vs. *MO-rbm8aATG* 1:200 p=0.5628 (not significant), *rbm8a^+/+^* vs. *rbm8a^Δ5/+^* p=0.605 (not significant), *rbm8a^+/+^* vs. *rbm8a^Δ3/ Δ3^* p=0.7655 (not significant), *rbm8a^+/+^* vs. *rbm8a^Δ3/ Δ5^* p=0.4289 (not significant); 6 dpf (**K**) wild type vs. control MO p=0.9567 (not significant), wild type vs. *MO-rbm8aATG* 1:125 p<0.0001, wild type vs. *MO-rbm8aATG* 1:150 p=0.5198 (not significant), wild type vs. *MO-rbm8aATG* 1:175 p=0.7147 (not significant), wild type vs. *MO-rbm8aATG* 1:200 p=0.288 (not significant), *rbm8a^+/+^* vs. *rbm8a^Δ5/+^* p=0.9462 (not significant), *rbm8a^+/+^* vs. *rbm8a^Δ3/ Δ3^* p=0.0208, *rbm8a^+/+^* vs. *rbm8a^Δ3/ Δ5^* p<0.0001 (see **Supplementary Data 2** for details). Scale bars in (**B**): 200 μm, applies to panels B,D,F,H; (**C**): 150 μm, applies to panels C,E,G,I.

We further tested the functionality of thrombocytes with reduced *rbm8a* function. Following laser-mediated injury of the dorsal aorta, *cd41*:EGFP-positive thrombocytes aggregated at the new wound site to occlude the damaged artery. The resulting time to occlusion (TTO) and number of aggregating thrombocytes provide a measure for a functional coagulation response ^74,75^. We did not observe any significant differences in TTO or number of aggregating thrombocytes between 6 dpf wild type and heterozygous *rbm8a^Δ5^* larvae (**Supplementary Fig. 2G,I, Supplementary Data 2**), homozygous *rbm8a^Δ3/Δ3^* or trans-heterozygous *rbm8a^Δ3/Δ5^* (**Supplementary Fig. 2H,J, Supplementary Data 2**).

Taken together, using our available genetic tools, these observations indicate that while *rbm8a*-perturbed larvae show reductions in thrombocyte numbers that are dependent on functional Rbm8a levels, the function of the thrombocytes is not affected.

### *rbm8a* perturbation impairs mRNAs encoding planar cell polarity components

To identify developmental mechanisms influencing thrombocyte formation and general hematopoiesis upon *rbm8a* perturbation, we sought to define the transcriptome of *rbm8a*-mutant zebrafish embryos at early developmental stages. Prior work has described the transcriptome of 21 hpf and 27 hpf *rbm8a^Δ5^*^/*Δ5*^ embryos, when mutants show clear signs of deterioration including onset of widespread apoptosis and necrosis^14^. We therefore analyzed the transcriptome of tailbud (9 hpf) *rbm8a^Δ5^*^/*Δ5*^ embryos as well as 24 hpf as comparison, capturing with the former the end of gastrulation when maternal Rbm8a protein and *rbm8a* mRNA levels start to wane (**Fig. 3 A-C**). The RNA-seq data can be browsed in our R/Shiny-based app RNA-seq Explorer with an interactive interface (http://imlspenticton.uzh.ch:3838/mosimann_p2452/; see Methods for details).

**Figure 3:**
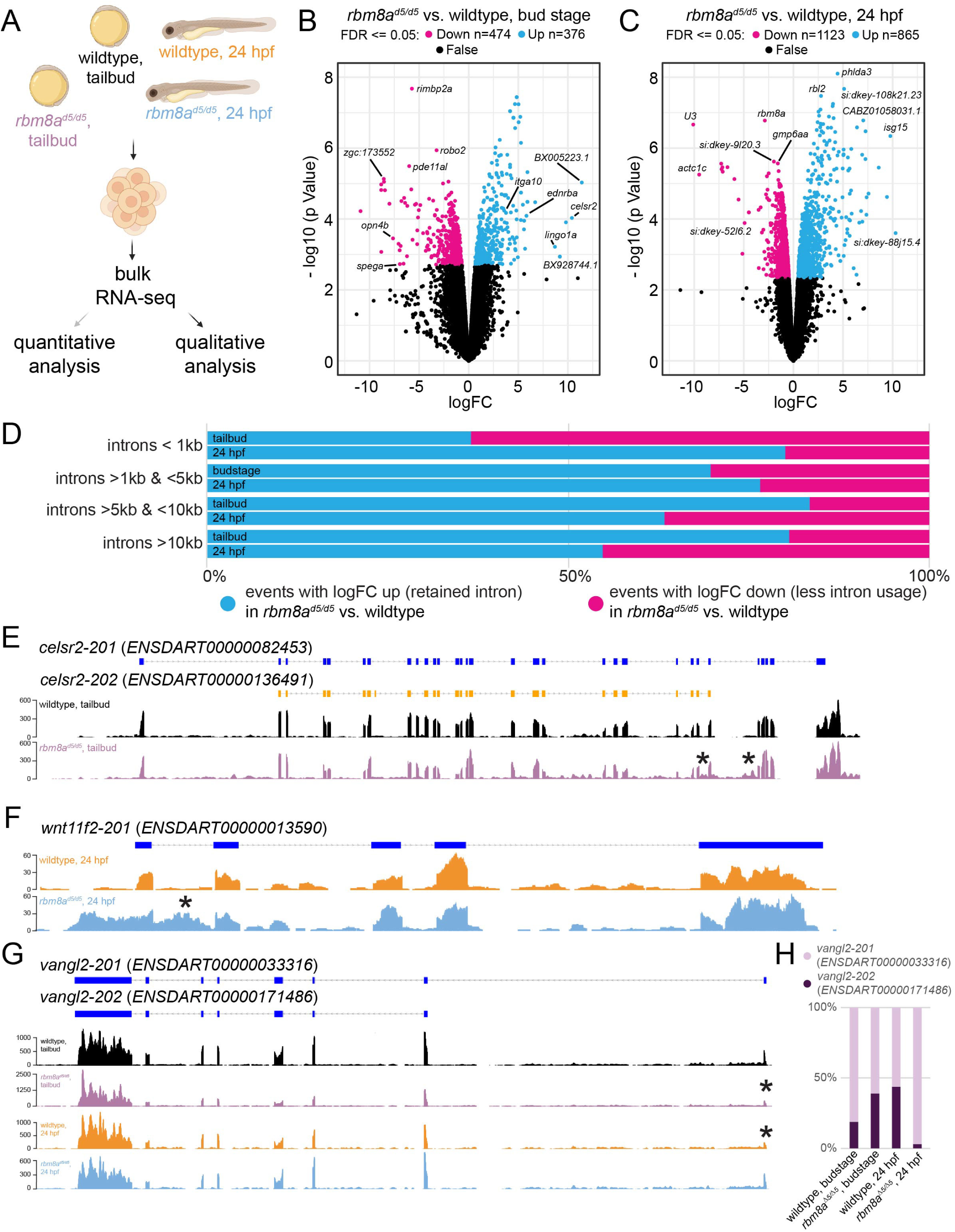
Intron retention in mRNAs encoding non-canonical Wnt/PCP components in zebrafish *rbm8a* mutants. (**A**) Experimental design of bulk RNA-seq experiment. Created with Biorender.com (Subscription: Individual, Agreement number: SN25WQAKB9). (**B**,**C**) Volcano plots of tailbud (**B**) and 24 hpf (**C**) comparisons between wild type versus *rbm8a*-mutant embryos; significantly down- (magenta) and up-regulated (blue) genes with named examples (see **Supplementary Data 3 and 4** for details). (**D**) Qualitative assessment of retained introns between wild type versus *rbm8a*-mutant embryos; note that retained introns are already apparent at tailbud stage (see **Supplementary Data 6** for details). (**E**, **F**) Read coverage plot of mRNA sequencing reads of non-canonical Wnt/PCP components *celsr2*, *wnt11f2* (former *wnt11*), and *vangl2* that feature mRNAs with retained introns (**E**,**F**) and/or differential transcript usage (**G**) in *rbm8a*-mutant embryos. (**H**) Differential transcript usage of *vangl2* transcripts at 24 hpf (exon 1 followed by long intron, asterisks).

Comparing *rbm8a^Δ5^*^/*Δ5*^ versus wild type embryos, we identified 850 differentially expressed genes (720 of which were protein coding) with FDR <= 0.05 at tailbud stage (**Fig. 3B, Supplementary Data 3**), and 1988 genes (1818 of which were protein-coding) with FDR <= 0.05 at 24 hpf (**Fig. 3C, Supplementary Data 4**). We used Metascape ^76^ to identify statistically enriched functional annotations of deregulated genes and their accumulative hypergeometric p-values (**Supplementary Fig. 3A,B, Supplementary Data 5**): similar to the reported transcriptomes of 21 and 27 hpf *rbm8a^Δ5^*^/*Δ5*^ embryos ^14^, at 24 hpf we observed enrichment for cell death and responses to stress among the upregulated genes, as well as downregulation of cell cycle processes and chromatin organization (**Supplementary Fig. 3A,B, Supplementary Data 5**). These transcriptional changes are in line with the progressive deterioration of *rbm8a^Δ5^*^/*Δ5*^ embryos as visible by the mutant and morphant phenotypes at that stage (**Fig. 1**) ^14^. In contrast, at tailbud stage, we observed broad deregulated gene categories associated with morphogenesis and general organ development (**Supplementary Fig. 3A,B, Supplementary Data 5**).

As part of the EJC, Rbm8a functions in NMD that removes mRNAs with retained introns or other splicing defects that occur routinely during native post-transcriptional processing ^12,13,77,78^. In *Drosophila*, loss of *rbm8a* causes selective retention of mRNAs with mis-spliced large introns due to defective NMD ^10,11^, and previous analysis of zebrafish *rbm8a* mutants documented stabilization of mRNAs with introns in their 3’ UTRs ^14^. Together, these data document both quantitative as well as qualitative impact on individual mRNAs upon EJC perturbation in different models. Given the depth of sequencing and early versus late timepoint in our dataset, we next asked if qualitative differences in individual transcripts could link Rbm8a deficiency with TAR Syndrome-associated processes.

We therefore scanned the tailbud stage and 24 hpf transcriptome of *rbm8a*-mutant embryos for differential exon and intron usage of individual mRNAs using DEXSeq (**Supplementary Fig. 3C**) ^79^. After filtering (retaining only protein-coding transcripts and further excluding U12 splice introns, transcripts with read counts below 100, adjusted p value per gene <= 0.05 and differential exon/intron usage event <= 0.1), we found 697 differential intron usage events in 564 genes (**Supplementary Data 6**) and 713 differential exon usage events in 502 genes (**Supplementary Data 7**) in *rbm8a^Δ5/Δ5^* mutants compared to wild type at tailbud stage (**Supplementary Fig. 3C**). In contrast, and again in line with progressive deterioration, we found 2220 differential intron usage events in 1594 genes (**Supplementary Data 6**) and 5439 differential exon usage events in 2316 genes (**Supplementary Data 7**) in *rbm8a^Δ5/Δ5^* compared to wild type at 24 hpf (**Supplementary Fig. 3C**). This analysis reveals a considerable qualitative difference in numerous mRNAs following *rbm8a* perturbation in zebrafish, including mRNAs with retained introns. Curiously, most of the retained introns were larger than 1 kb at tailbud stage, compared to retained introns of all sizes at 24 hpf when the *rbm8a^Δ5^*^/*Δ5*^ embryos were declining (**Fig. 3D**). We speculate that at tailbud stage, given Rbm8a involvement in the EJC, problems with longer introns start to accumulate while smaller introns seem to still being spliced properly; at 24 hpf, introns of all sizes are affected.

We again used Metascape ^76^ to identify statistically enriched functional annotations of the genes with differential intron usage events at tailbud stage and 24 hpf (**Supplementary Fig. 3C**). We observed statistically enriched annotations for cell-cell adhesion and cell junction organization at tailbud stage and at 24 hpf stage amongst others (**Supplementary Fig. 3E, Supplementary Data 8**). When we looked manually through the genes associated with the enriched GO terms, we found *celsr2*, *wnt11*, and *prickle1*, as well as several additional mRNAs encoding components of the non-canonical Wnt/PCP signaling, including *wnt5b* and *vangl2* (**Fig. 3E-G, Supplementary Fig. 3F**), among irregular transcripts with developmental contributions at both analyzed stages. While per *se not* enough to evoke a GO term involving non-canonical Wnt signaling, even though the GO terms cell-cell adhesion and cell junction organizations are cellular processes linked to PCP signaling, significant intron retention of these genes indicates possibly perturbed non-canonical Wnt pathway involved in PCP signaling. PCP signaling equips tissues with a polarity axis resulting in collective morphogenetic events, such as the orientation of subcellular structures and cell rearrangements ^49^. These mRNAs showed variable, yet considerable sequencing reads in individual introns, indicating seemingly minor but significant accumulation of mis-spliced transcripts with retained introns (**Fig. 3E-G, Supplementary Fig. 3F**). Such intron retention is predicted to lead to unproductive protein translation or truncated, functionally perturbed polypeptides ^12,13^, potentially lowering the effective dose of individual non-canonical Wnt/PCP components. We also noted that one transcript isoform of *celsr2*, encoding one of several Flamingo family receptors for non-canonical Wnt/PCP ligands ^49,80^, was among the top-upregulated mRNAs at tailbud stage in *rbm8a^Δ5^*^/*Δ5*^ embryos (logFC 10.4, FDR=0.013); at 24 hpf, *celsr2* mRNAs showed selective intron retention of intron 15 (logFC 2.4, p=0.00015) (**Fig. 3B,C, Supplementary Fig. 3F**). Transcripts of the key PCP component *vangl2* showed differential variant use between our sequenced timepoints (**Fig. 3G,H**).

Taken together, our analysis indicates that *rbm8a*-mutant embryos already show developmental anomalies at the bulk transcriptome level at the end of gastrulation, when maternal *rbm8a* mRNA and Rbm8a protein contribution is starting to fade. From our qualitative assessment of mRNAs at tailbud and 24 hpf stage, we hypothesized that *rbm8a*-perturbed zebrafish embryos feature a mild, yet functionally significant attenuation of non-canonical Wnt/PCP signaling involved in convergence and extension movements and other cell polarity coordination. Defective PCP signaling could impact the proper migration, and subsequently influence the cell fate determination of, the LPM stripes and the hematopoietic/endothelial progenitors.

### Zebrafish Rbm8a deficiency causes aberrant LPM morphology

As the phenotypes in TAR Syndrome arise in LPM-derived organs and cell types, we next sought to determine if the early LPM develops normally upon *rbm8a* perturbation. To observe LPM formation and patterning after gastrulation, we performed light sheet-based live imaging using the transgenic reporter *scl:EGFP* that marks the emerging hematopoietic and endothelial progenitors in the medial LPM from early somitogenesis stages (**Fig. 4A**) ^19,81,82^. In wild type, the bilateral *scl*-expressing LPM stripes gradually widen and thicken before starting an anterior-to-posterior midline migration around the 5 somite-stage (**Fig. 4B,E,F**). In contrast, *rbm8a* knockdown with *MO-rbm8aATG* resulted in reduced area and volume of the *scl:EGFP*-expressing LPM in the same developmental timespan (**Fig. 4C,E,F**). At the imaging endpoint at 10-11 somites, EGFP fluorescence from *scl:EGFP* showed no discernible difference in intensity between wild type and morphants, yet their overall morphology in mutants appeared wider apart and less converged to the midline (**Fig. 4B,C,E,F**). This reduced convergence is unlikely due to developmental delay, as we stage-matched the analyzed embryos by morphology and not absolute time post-fertilization.

**Figure 4:**
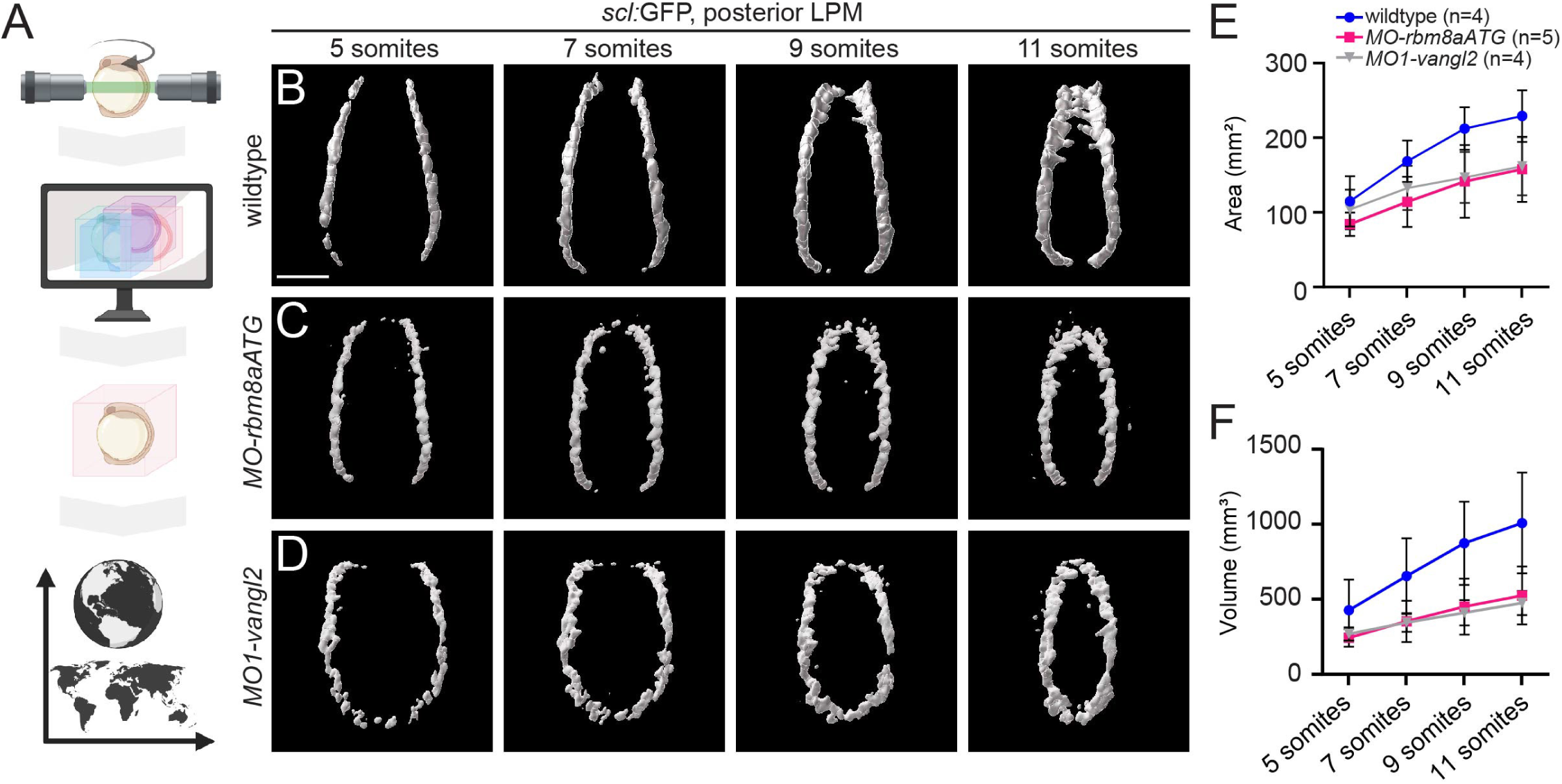
*rbm8a* and *vangl2* perturbations reduce the rate of posterior LPM growth. (**A**) Schematic of the workflow to image the zebrafish using Light sheet microscope from 4 different angles, assemble in a 3D image (represented by the globe) and measure volume and area of the PLPM *scl:GFP* surface rendering (represented by the map). Created with Biorender.com (Subscription: Individual, Agreement number: IG25WQA6FD). (**B**-**D**) Surface rendering of GFP signal from light sheet-based timelapse imaging of *scl:GFP* transgenic zebrafish embryos. Wild type (**A**), *rbm8a* morphant (**B**), and *vangl2* morphant (**C**) embryos at 5, 7, 9 and 11 somite stages, dorsal view, anterior to the top, posterior end of the LPM at bottom. (**D,E**) Area and volume of *scl*:GFP-expressing LPM territory measured with Imaris, comparing wild type (n=4), *rbm8a* morphant (n=5) and *vangl2* morphant (n=4) measurements depicted in color groups corresponding to sample groups. Note how area in wild type increases by x˄2 (**D**) and volume by x˄3 (**E**), revealing reduced growth rates of the posterior *scl*:GFP territory upon *rbm8a* and *vangl2* perturbations. Datapoints shown are average with standard deviation, significance calculated by 2-way Anova. Area (p<0.0001) and volume (p<0.0001) are significantly different in *MO-rbm8aATG*- and *MO1-vangl2*-injected embryos compared to stage-matched wild type embryos at the analyzed timepoints (see **Supplementary Data 9** for details). Scale bar in (**B**): 150 μm, applies to all panels in B-D.

Reflective of the central role of Vangl2 in relaying PCP signaling ^83–85^, zebrafish embryos mutant for *vangl2* and morphants show severe convergence and extension defects, resulting in embryos with shortened, wider posterior trunks and compacted tails ^60,84,86–91^. Consistent with these defects, *vangl2* MO knockdown resulted in reduced volume and width of the *scl:EGFP*-expressing LPM stripes with no discernible impact on fluorescent reporter levels (**Fig. 4D-F, Supplementary Data 9**). Together, these observations indicate that the *scl*-expressing LPM in both *rbm8a*- and in *vangl2*-perturbed embryos develops an aberrant morphology.

### *rbm8a*-mutant embryos show convergence and extension defects

Non-canonical Wnt/PCP signaling controls cell migration during convergence and extension movements that drive zebrafish embryonic axis elongation ^83,85,89,91–93^. Hallmarks of defective convergence and extension include reduced length of the anteroposterior axis and increased width of the neural plate and somites ^51,60,61,89,90,93–96^. To document and quantify if *rbm8a* perturbation causes convergence and extension defects in zebrafish, we measured morphometric parameters during early somite stages of wild type and *rbm8a*-mutant embryos: measured parameters included axis length, somite and neural plate width, three morphometric parameters affected by reduced convergence and extension in PCP mutants. To assign landmarks for these measurements, we combined *in situ* hybridization for *dlx3b*, *tbxta*, *myoD*, and *hgg1* as markers of the neural plate border, chordamesoderm, somites, and prechordal plate, respectively ^97–101^.

Compared to stage-matched wild type siblings, *rbm8a^Δ5/Δ5^* mutants as well as *rbm8a* morphants showed significantly reduced axis lengths concomitant with a decrease in axis angle at the analyzed 6-11 somite stages (**Fig. 5A, Supplementary Data 10**). The axis length and angle discrepancy was already detectable at the first developmental timepoint we measured. While *rbm8a^Δ5/Δ5^* mutant and *rbm8a* morphant embryos continued to extend in length, they fell short of reaching wild type lengths at the end of our stage series (**Fig. 5B,C**). In contrast, *rbm8a^Δ3/Δ5^* and *rbm8a^Δ3/Δ3^* mutant embryos did not show significant changes in axis length or axis angle at the 6-7 somite timepoint, but continued to being slightly shorter compared to wild type towards the end of our measurements (**Fig. 5B,C**). Further, compared to wild type embryos, *rbm8a^Δ5/Δ5^* mutant embryos featured wider neural plates (**Fig. 5D**) throughout our timecourse, while the *rbm8a* morphants, as well as *rbm8a^Δ3/Δ5^* and *rbm8a^Δ3/Δ3^* mutant embryos did not, despite a slight increase at the 8-9 somite stage (**Fig. 5D**). Lastly, *rbm8a^Δ5/Δ5^* mutant embryos had significantly wider somites throughout the timecourse compared to wild type embryos (**Fig. 5E**). The somite width of the *rbm8a* morphants was significantly reduced as well compared to wild type (except at the 8-9 somite stage), however their somites were not as wide as the somites of the *rbm8a^Δ5/Δ5^* mutants (**Fig. 5E**). In *rbm8a^Δ3/Δ5^* and *rbm8a^Δ3/Δ3^* mutants, the somite width was unchanged compared to wild type embryos at all timepoints (**Fig. 5E**).

**Figure 5:**
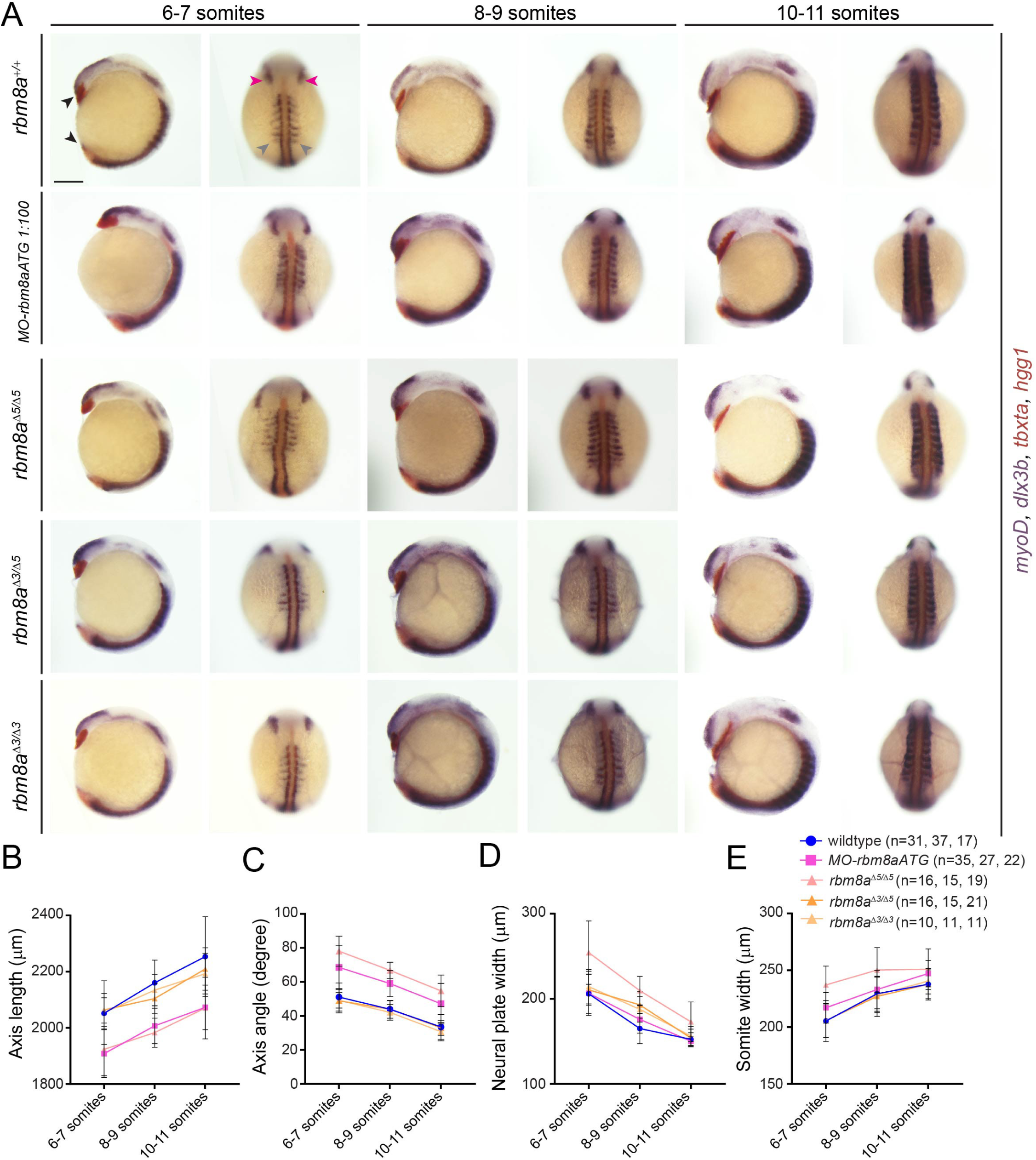
*rbm8a* perturbation results in convergence and extension defects. (**A**) mRNA *in situ* hybridization for *myoD*, *dlx3b*, *tbxta*, and *hgg1* as landmarks to measure morphometric parameters associated with convergence and extension as influenced by non-canonical Wnt/PCP signaling. Zebrafish embryos are shown as lateral views (odd columns, anterior to the top, ventral to the left) and dorsal views (even columns, anterior to the top). Black arrowheads depict anterior and posterior extent of the body axis in lateral views, pink and grey arrowheads depict otic placodes and somites at measured position in dorsal views. Axis length and axis angle (marked by black arrowheads) were measured from the odd panels. Neural plate (between the otic placodes at beginning of notochord, pink arrowheads) and somite width (last 3 somites, grey arrowheads) were measured from the even panels. (**B-E**) Morphometric measurements for axis length (**B**), axis angle (**C**), neural plate width (**D**), and somite width (**E**) from individual embryo (wild type, mutant, morphant) images at different developmental stages. Datapoints shown are average with standard deviation, significance calculated by Mann Whitney test: axis length (**B**) 6-7 somite stage wild type vs. *MO-rbm8aATG* p<0.0001, wild type vs. *rbm8a^Δ5/Δ5^* p<0.0001, wild type vs. *rbm8a^Δ3/Δ5^* p=0.4394 (not significant), wild type vs. *rbm8a^Δ3/Δ3^* p=0.9347 (not significant), 8-9 somite stage wild type vs. *MO-rbm8aATG* p<0.0001, wild type vs. *rbm8a^Δ5/Δ5^* p<0.0001, wild type vs. *rbm8a^Δ3/Δ5^* p=0.0048, wild type vs. *rbm8a^Δ3/Δ3^* p=0.1643 (not significant), 10-11 somite stage wild type vs. *MO-rbm8aATG* p=0.0001, wild type vs. *rbm8a^Δ5/Δ5^* p=0.0003, wild type vs. *rbm8a^Δ3/Δ5^* p=0.0945 (not significant), wild type vs. *rbm8a^Δ3/Δ3^* p=0.1219 (not significant); axis angle (**C**) 6-7 somite stage wild type vs. *MO-rbm8aATG* p<0.0001, wild type vs. *rbm8a^Δ5/Δ5^* p<0.0001, wild type vs. *rbm8a^Δ3/Δ5^* p=0.3325 (not significant), wild type vs. *rbm8a^Δ3/Δ3^* p=0.4449 (not significant), 8-9 somite stage wild type vs. *MO-rbm8aATG* p<0.0001, wild type vs. *rbm8a^Δ5/Δ5^* p<0.0001, wild type vs. *rbm8a^Δ3/Δ5^* p=0.7492 (not significant), wild type vs. *rbm8a^Δ3/Δ3^* p=0.2633 (not significant), 10-11 somite stage wild type vs. *MO-rbm8aATG* p<0.0001, wild type vs. *rbm8a^Δ5/Δ5^* p<0.0001, wild type vs. *rbm8a^Δ3/Δ5^* p=0.439 (not significant), wild type vs. *rbm8a^Δ3/Δ3^* p=0.342 (not significant), neural plate width (**D**) 6-7 somite stage wild type vs. *MO-rbm8aATG* p=9978 (not significant), wild type vs. *rbm8a^Δ5/Δ5^* p<0.0001, wild type vs. *rbm8a^Δ3/Δ5^* p=0.4437 (not significant), wild type vs. *rbm8a^Δ3/Δ3^* p=0.3001 (not significant), 8-9 somite stage wild type vs. *MO-rbm8aATG* p=0.0086, wild type vs. *rbm8a^Δ5/Δ5^* p<0.0001, wild type vs. *rbm8a^Δ3/Δ5^* p<0.0001, wild type vs. *rbm8a^Δ3/Δ3^* p=0.0008, 10-11 somite stage wild type vs. *MO-rbm8aATG* p=0.8324 (not significant), wild type vs. *rbm8a^Δ5/Δ5^* p=0.0198, wild type vs. *rbm8a^Δ3/Δ5^* p=0.5477 (not significant), wild type vs. *rbm8a^Δ3/Δ3^* p=0.5448 (not significant); somite width (**E**) 6-7 somite stage wild type vs. *MO-rbm8aATG* p<0.0001, wild type vs. *rbm8a^Δ5/Δ5^* p<0.0001, wild type vs. *rbm8a^Δ3/Δ5^* p=0.2296 (not significant), wild type vs. *rbm8a^Δ3/Δ3^* p=0.7951 (not significant), 8-9 somite stage wild type vs. *MO-rbm8aATG* p=0.1853 (not significant), wild type vs. *rbm8a^Δ5/Δ5^* p<0.0001, wild type vs. *rbm8a^Δ3/Δ5^* p=0.2642 (not significant), wild type vs. *rbm8a^Δ3/Δ3^* p=0.6899 (not significant), 10-11 somite stage wild type vs. *MO-rbm8aATG* p=0.0028, wild type vs. *rbm8a^Δ5/Δ5^* p=0.0003, wild type vs. *rbm8a^Δ3/Δ5^* p=0.5769 (not significant), wild type vs. *rbm8a^Δ3/Δ3^* p=0.4317 (not significant) (see **Supplementary Data 10** for details). Scale bar in (**A**): 250 μm, applies to all panels in A.

Together, while milder than phenotypes observed in mutants for core regulators of the process such as *kny* or *tri*/*vangl2* ^89,96^, our results indicate that somite-stage *rbm8a*-deficient zebrafish embryos show phenotypes consistent with convergence and extension defects.

### Non-canonical Wnt/PCP signaling is sensitive to Rbm8a levels

We next tested whether non-canonical Wnt/PCP signaling is sensitive to attenuated Rbm8a levels. We used previously established morpholino oligonucleotides to reduce the levels of the intron-retained PCP ligands, *wnt5b* and *wnt11f2*, as well as transmembrane PCP core components *fzd7a* and *vangl2*: each morpholino has a well-established injection dose that recapitulates the recessive mutant phenotypes of each respective gene as previously validated per current guidelines ^52,68,102,103^.

To determine whether reduction of non-canonical PCP core components sensitizes zebrafish embryos to reduced Rbm8a levels, we co-injected *MO-rbm8aATG* (**Fig. 6A,B**) together with each PCP core component-targeting morpholino at sub-threshold concentrations into wild type embryos **(Fig. 6C-F**). We used morphology of the body axis as a read-out to score the level of interaction between *rbm8a* and PCP core components at 48 hpf. Combined knockdown of individual PCP core component genes and *rbm8a* resulted in phenotypes that approached the full-dose knockdown of each Wnt/PCP gene (**Fig 6C-F**). For *fzd7a*, *vangl2*, and *wnt11f2*, over 90% of all co-injected embryos became phenotypic, indicating a strong genetic interaction with *rbm8a* resulting in a predominant severe phenotype class and lower number of moderate phenotypic embryos (**Fig. 6G, Supplementary Data 11**). *wnt5b* attenuation showed the mildest, yet still significant, response to *rbm8a* co-attenuation with 40% embryos becoming phenotypic (**Fig. 6E,G**). The combined attenuation resulted in embryos with shortened body axes and concomitant circulation defects, as common to perturbations of PCP signaling that affects embryo elongation by convergence and extension ^49,51,89–91,93^.

**Figure 6:**
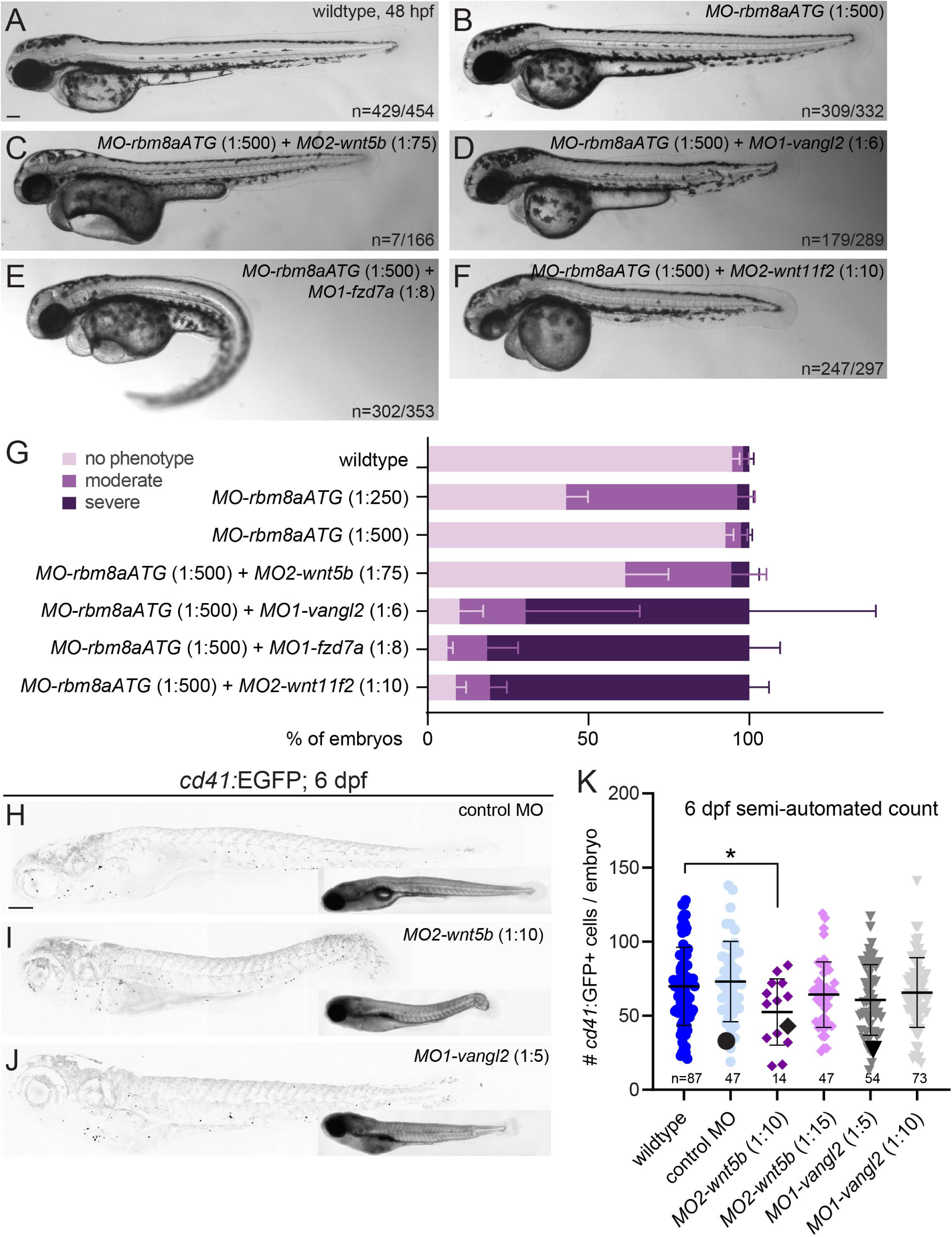
*rbm8a* attenuation sensitizes zebrafish to non-canonical Wnt/PCP defects. (**A**-**F**) 2 dpf lateral views of representative zebrafish larvae analyzed for each genetic perturbation. Anterior to the left. (A) Uninjected wild type as phenotype reference. (**B**) Suboptimal (1:500) *MO-rbm8aATG* injection that does not result in a visible phenotype (as compared to higher doses) and provides a baseline for *rbm8a* attenuation. (**C-F**) Representative phenotypes following sequential injection of suboptimal *MO-rbm8aATG* and suboptimal doses of individual, validated morpholinos against the non-canonical Wnt/PCP components *fzd7* (**C**), *vangl2* (**D**), *wnt5b* (**E**), and *wnt11f2* (**F**). (**G**) Observed phenotypes in percent of embryos. (**H-K**) Reduced function of *vangl2* and *wnt5b* results in reduced *cd41*:GFP-expressing thrombocytes at 6 dpf. (**H**-**J**) Lateral, inverted greyscale views of 6 dpf *cd41:GFP*-transgenic embryos confocal-imaged for GFP fluorescence, inserts depict brightfield view of imaged embryos and overall phenotype reference. Note how suboptimal dosing retains longer viability and less morphological impact (compare to **D**,**E**). (**K**) Thrombocyte quantification at 6 dpf reveals mild, yet dose-dependent reduction in cell number upon *vangl2* and *wnt5b* perturbation. Individual datapoints (total number of *cd41:GFP*-positive cells per embryo) shown with mean and standard deviation, significance calculated by Mann-Whitney test: 6 dpf (**K**) wild type vs. control MO p=0.4981 (not significant), wild type vs. *MO2-wnt5b* 1:10 p=0.0407, wild type vs. *MO2-wnt5b* 1:15 p=0.2364 (not significant), wild type vs. *MO1-vangl2 1:5* p=0.0533 (not significant), wild type vs. *MO1-vangl2* 1:10 p=0.3828 (not significant) (see **Supplementary Data for 11** for details). Scale bar in (**A**): 100 μm, applies to panels A-F; (**H**): 150 μm, applies to panels H-J.

To deduce if reducing PCP signaling has a downstream impact on thrombocyte formation, we revisited our *cd41:EGFP*-based thrombocyte counts (**Fig. 2**). We injected cohorts of *cd41:EGFP* embryos with two sub-optimal doses of morpholinos against *vangl2* and *wnt5b*, respectively; injection of each resulted in viable larvae with shortened body axis while establishing seemingly normal blood circulation in the observed timeframe (**Fig. 6H-K**). Compared to 6 dpf wild type and control morpholino-injected larvae, both *vangl2*- and *wnt5b*-attenuated zebrafish harbored reduced numbers of *cd41:EGFP*-expressing thrombocytes with a clear dose response (**Fig. 6K, Supplementary Data 2**), and a significant decrease in *MO2-wnt5b* (1:10)-injected larvae.

Taken together, our injection-based genetic interaction series established that reduced levels of *rbm8a* sensitize zebrafish embryos to perturbations in components of the non-canonical Wnt/PCP pathway during early development, with consequences to later stages. These observations connect with our morphometric data (**Fig. 4,5**), indicating that *rbm8a*-mutant zebrafish show features of defective convergence and extension that also affect the hematopoietic progenitors among the LPM.

### Perturbing *rbm8a* or *vangl2* impacts hematopoietic gene expression

Following bilateral stripe emergence, *scl:EGFP*-positive cells in the converging *drl:mCherry*-positive LPM form hematopoietic, and endothelial fates in the trunk (**Fig. 7A**) ^20,82^. We therefore sought to evaluate possible changes to hematopoietic and endothelial progenitor formation and patterning upon *rbm8a* and *vangl2* perturbation, respectively. If attenuation of *rbm8a* results in PCP signaling defects that subsequently cause LPM migration and hematopoietic fate disruptions, we would also anticipate that interrupted PCP signaling in *vangl2* mutants reveals comparable phenotypes. We performed mRNA *in situ* hybridization (ISH) on *rbm8a^Δ5/Δ5^*, *rbm8a^Δ3/Δ3^*, *MO-rbm8aATG*-injected, *vangl2^vu^*^67^*^/vu67^*, and wild type control embryos with a panel of marker genes expressed in in hematopoietic and endothelial progenitors (*sox7*, *kdrl*, *gata1*, *gfi1aa*, *gfi1b*, and *runx1*) (**Fig. 7B-H**). *In situ* hybridization for the pan-trunk muscle marker *myoD* provided a reference across conditions and for overall embryo morphology (**Fig. 7B**). We stage-matched embryos to wild type and control *MO*-injected controls by somite number and overall morphology to ensure valid comparisons and quantifications.

**Figure 7:**
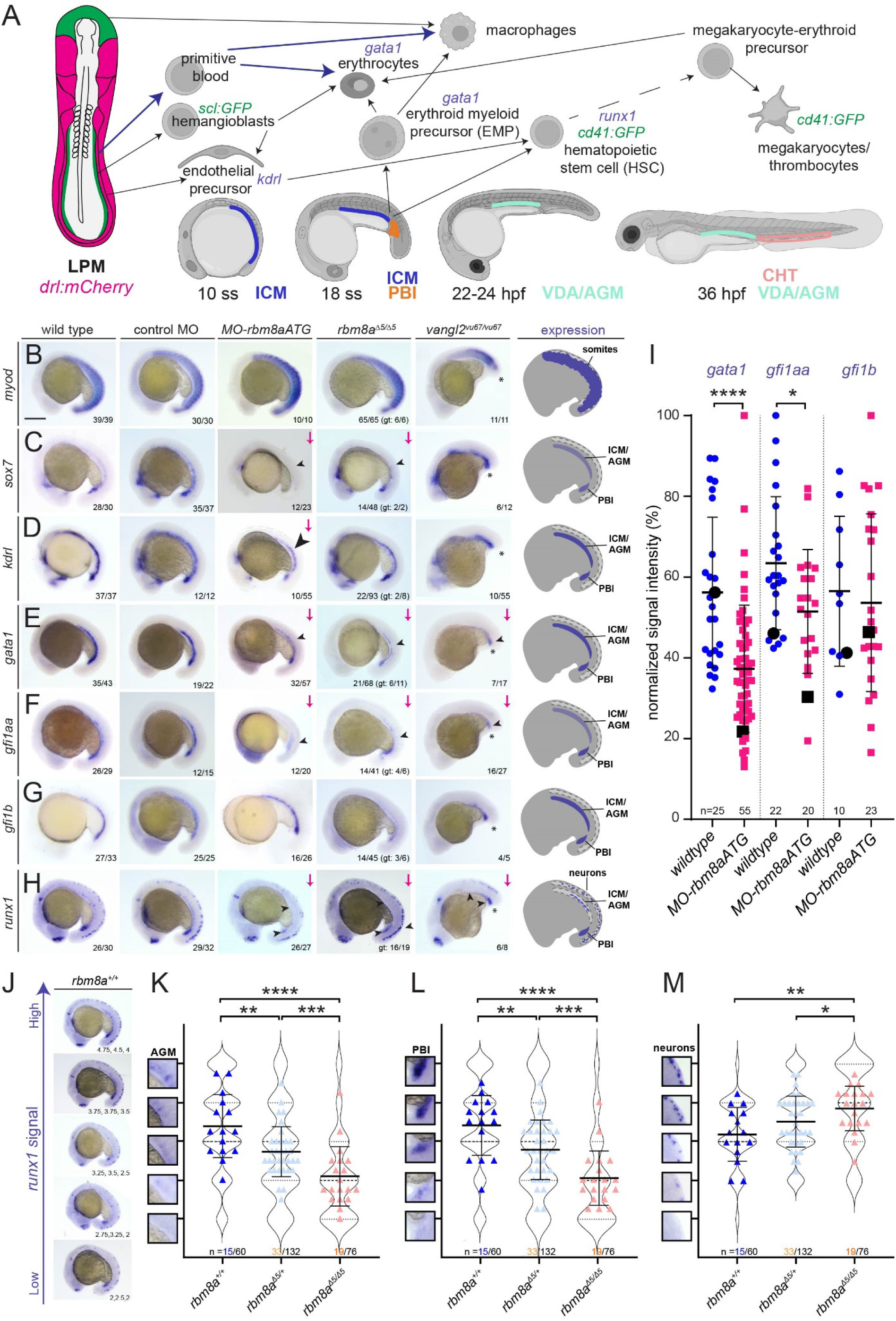
Select changes in hematopoietic marker expression in *rbm8a*- and *vangl2*-mutant zebrafish embryos. (**A**) Schematic of basic hematopoietic lineage and marker relationships. Created with Biorender.com (Subscription: Individual, Agreement number: IZ25WQERBS). AGM: aorta-gonadal-mesonephros area; ICM: intermediate cell mass; PBI: posterior blood island; CHT: caudal hematopoietic territory; VDA: ventral dorsal aorta. (**B**-**F**) Expression patterns for individual marker genes in wild type, morpholino-injected, *rbm8a*- and *vangl2*-mutant zebrafish embryos at stage-matched 18 somites. Schematic of zebrafish embryo in the last panel in **B-H** shows where marker expression is expected and analyzed. (**B**) *myoD* as marker for somitic muscles shows comparable expression across conditions, with *vangl2* mutants depicting the short, deteriorating tail typical for these mutants (asterisk). (**C**) The early endothelial marker *sox7* is reduced in the trunk upon *rbm8a* perturbation (arrowheads) and seemingly normal in *vangl2* mutants (asterisk). (**D**) The endothelial marker *kdrl* is decreased similarly as *sox7*. (**E**) The erythroid marker *gata1* is reduced in all *rbm8a*-perturbed embryo conditions (arrowheads) and only retained in posterior-most LPM cells of *vangl2* mutants (asterisk). (**F**) The hematopoietic progenitor marker *gfi1aa* is reduced in all *rbm8a*-perturbed embryo conditions (arrowheads) and barely detectable in *vangl2*-mutant embryos (asterisk). (**G**) In contrast, *gfi1b* is unchanged in all conditions. (**H**) The hematopoietic progenitor marker *runx1* in *rbm8a* morphants and mutants is reduced or absent in the trunk and posterior blood island (PBI) (embryo midline, future dorsal aorta, arrowheads); *vangl2* mutants also show reduced trunk expression (arrowhead) yet retain PBI *runx1* expression (asterisk). (**I**) mRNA *in situ* hybridization-based quantification n of *gata1*, *gfi1aa*, and *gfi1b* expression levels; individual data points (normalized signal intensity/embryo) shown with mean and standard deviation, significance calculated by Mann-Whitney test: *gata1* ISH signal intensity in wild type vs. *MO-rbm8aATG* p<0.0001, *gfi1aa* ISH signal intensity in wild type vs. *MO-rbm8aATG* p=0,031, gfi1b ISH signal intensity in wild type vs. *MO-rbm8aATG* p=0.7727 (not significant). (**J-M**) mRNA *in situ* hybridization-based quantification of *runx1* in select areas (aorta-gonadal-mesonephros area (AGM): embryo midline, future dorsal aorta), posterior blood island (PBI), neurons). (**J**) Representative scale of *runx1* signal ranked from low to high in *rbm8a* wild type siblings. The numbers on the bottom represent the average scores for AGM, PBI and neurons, respectively. (**K**) Signal in AGM. (**L**) Signal in PBI. (**M**) Signal in dorsal neurons. Individual data points (average of 4 scores per embryo) shown with mean and standard deviation, significance calculated by Mann-Whitney test: *runx1a* signal intensity in AGM (**K**) in wild type vs. *rbm8a^Δ5/+^* p=0.0048, wild type vs. *rbm8a^Δ5/Δ5^* p<0.0001, *rbm8a^Δ5/+^* vs. *rbm8a^Δ5/Δ5^* p=0.0009; PBI (**L**) in wild type vs. *rbm8a^Δ5/+^* p=0.0077, wild type vs. *rbm8a^Δ5/Δ5^* p<0.0001, *rbm8a^Δ5/+^* vs. *rbm8a^Δ5/Δ5^* p=0.0007; neurons (**M**) wild type vs. *rbm8a^Δ5/+^* p=0.1604 (not significant), wild type vs. *rbm8a^Δ5/Δ5^* p=0.005, *rbm8a^Δ5/+^* vs. *rbm8a^Δ5/Δ5^* p=0.0481. Violin plot in background depicts population of the not averaged scores. Numbers (n)=average/total (see **Supplementary Data 12** for details). Scale bar in (**B**): 100 μm, applies to all panels in B-H.

Compared to wild type and control MO-injected embryos, we observed decreased and patchy expression of the early pan-endothelial cell markers *sox7* (**Fig. 7C**) and *kdrl* (**Fig. 7D**) in the trunk of *rbm8a^Δ5/Δ5^* genotyped embryos (n=2/2 *sox7*, total n=14/48; n=2/8 *kdrl*, total n=22/93), as well as in *MO-rbm8aATG*-injected embryos (n=12/23 *sox7*, n=10/55 *kdrl*) ^19,104,105^. In comparison, the anterior endothelium and endocardial progenitor fields showed variable reduction of *sox7* and *kdrl* expression in *MO-rbm8aATG*-injected embryos (**Fig. 7C,D**). While still expressing *sox7* and *kdrl* at overall comparable levels to wild type siblings, *vangl2^vu76/vu67^* embryos (n=6/12 *sox7*, n=2/10 *kdrl*) showed no discernible midline fusion of the bilateral *kdrl*-expressing progenitors in the trunk at 18 somite stage (ss), in line with perturbed convergence and extension and midline migration **(Fig. 7C,D**).

The erythroid progenitor marker *gata1* at 18 ss reveals the emerging primitive red blood cells in the intermediate cell mass of the trunk (**Fig. 7A,E**) ^19,106^. In both stage-matched, genotyped *rbm8a^Δ5/Δ5^* (n=6/11, total n=47/68) and *MO-rbm8aATG*-injected embryos (n=32/57), *gata1* expression was significantly reduced or patchy compared to wild type or control MO-injected embryos by visual inspection (**Fig. 7E**) and *in situ hybridization* signal quantification (wild type n=25, *MO-rbm8aATG*-injected n=24) (**Fig. 7I, Supplementary Data 12**). Similarly, *gata1* expression in *vangl2^vu76/vu67^* embryos remained in the posterior-most LPM, yet was greatly reduced in the trunk (n=7/17) (**Fig. 7E**).

The transcription factors GFI1 and GFI1B are key determinants of myeloid fate potential towards erythroid/megakaryocyte and lymphoid fates in mammals, and mutations in human *GFI1B* cause thrombocyte deficiencies ^37^. In zebrafish, the orthologs *gfi1aa* and *gfi1b* influence primitive erythroid precursor formation and are expressed in intermediate and definitive precursors (**Fig. 7A**) ^40,41,107^. Compared to wild type and control MO-injected embryos, both genotyped *rbm8a^Δ5/Δ5^* (n=4/6, total n=27/41) and *MO-rbm8aATG*-injected embryos (n=12/20) displayed significantly reduced *gfi1aa* expression, which we further verified by signal intensity comparison (wild type n=23, *MO-rbm8aATG*-injected n=21). (**Fig. 7F,I**). *gfi1aa* expression was also markedly reduced in *vangl2^vu76/vu67^* embryos compared to wild type siblings (n=16/27) (**Fig. 7F**). In contrast, *gfi1b* expression showed no significant quantitative differences (**Fig. 7I**), yet we observed individual genotyped *rbm8a^Δ5/Δ5^* embryos with patchy *gfi1b* expression (n=3/6, total 14/45) (**Fig. 7G**).

Lastly, *runx1* expression that broadly marks prospective hematopoietic progenitors including future stem cell precursors in the trunk at 18 ss (**Fig. 7A**) ^108^ was reduced in genotyped *rbm8a^Δ5/Δ5^* (n=16/19) and *MO-rbm8aATG*-injected embryos (n=26/27) (**Fig. 7H**). We further verified our observations by signal intensity comparison of *runx1*-positive cells in the aorta-gonadal-mesonephros area (AGM) and posterior blood island (PBI), which showed a significant decrease in *runx1*-positive cells in those areas in both *rbm8a^Δ5/+^* and *rbm8a^Δ5/Δ5^* mutant embryos compared to wild type siblings (**Fig. 7J-L**). Notably, the spinal cord neurons and other domains expressing *runx1* seemed unaffected in number (**Fig. 7F**), a hallmark of mutants with selective hematopoietic defects ^109^, but showed slightly higher expression levels in our quantitative signal intensity comparison (**Fig. 7M, Supplementary Data 12**). Although *vangl2^vu67/vu67^* embryos maintained *runx1* expression in the eye, ear, and dorsal spinal cord neurons, they displayed selectively reduced or absent *runx1* expression in the presumptive hematopoietic stem cell progenitors at the midline as well as in the PBI (n=6/8) (**Fig. 7H**). This impaired *runx1* expression in both *rbm8a^Δ5/Δ5^* and *vangl2^vu76/vu67^* embryos is unlikely to be caused by developmental delay or the severely perturbed midline migration in *vangl2^vu76/vu67^* embryos, as *runx1* expression is stronger in earlier bilateral LPM progenitors before confining to hematopoietic precursors in the trunk (**Fig. 7A,H**) ^108,109^.

While only including a limited sampling of perturbations and timepoints, together these data document select changes in hematopoietic progenitor gene expression upon *rbm8a* and *vangl2* defects. Our data proposes a model in which attenuated Rbm8a levels and concomitant EJC reduction result in retention of mis-spliced mRNAs encoding non-canonical Wnt/PCP genes (among others), lowering the effective protein concentration of several PCP pathway components; this results in sub-optimal convergence and extension and cell migration including LPM progenitors, with subsequent impact on hematopoietic progenitor numbers and/or gene expression. In line with this model, zygotic *rbm8a*-mutant and -knockdown zebrafish feature select and significant blood and endothelial lineage defects already prior to the onset of heartbeat and circulation.

## DISCUSSION

The striking phenotype combination of TAR syndrome – affecting selective blood and limb skeleton features as well as heart, kidney, and craniofacial structures – has remained challenging to consolidate with a unifying disease mechanism. The identification of hypomorphic defects in the ubiquitous EJC factor Rbm8a in TAR syndrome patients has further complicated possible explanations of how the selective phenotypes arise. Investigating the developmental impact following different means of reducing Rbm8a protein levels in zebrafish, we propose that TAR syndrome involves early embryonic perturbations in LPM formation and patterning via attenuated PCP signaling. Our work links sensitizing reduction of pathways essential for embryo morphogenesis to retained introns in individual mRNAs upon Rbm8a reduction, proposing a developmental mechanism to consolidate the seemingly divergent phenotypes of TAR syndrome as LPM-affecting birth defect.

Disruption of a molecular mechanism that is shared in the development of diverse cell fates by means of clonal cell relationship, evolutionary co-option, or signaling interactions between progenitors can lead to complex syndromic phenotypes. Mutations in pleiotropic factors such as ribosomal or DNA repair components have been linked to hematological disorders with additional structural defects that include the limbs as observed in Diamond-Blackfan Anemia and Fanconi’s Anemia, respectively ^110–115^. Curiously, distinct rare congenital cases with combined amegakaryocytic thrombocytopenia and radial-ulnar anomalies have been reported (CTRUS or RUSAT, OMIM # 605432) ^116^. Mutations in *HOXA11* have been linked to thrombocytopenia with concurrent proximal fusion of radius and ulna in affected patients ^117^, and *Hox* gene mutations are hypothesized to underlie similar phenotypic manifestations in rare cases ^113^. Individual patients with initial platelet deficiency that improves with age and deficient pronation-supination of the forearm carry mutations in *MECOM1* encoding the transcription factor EVI-1^118^. Both Hoxa11 and Evi-1 have been associated with hematological, forelimb, and other LPM-linked disorders ^119–128^. Independent of underlying molecular mechanism, establishing a developmental connection between affected cell types in syndromic congenital disease provides a potent framework to expand diagnosis, phenotype assessment, and long-term care of affected patients. The multi-lineage potential of early LPM provides a developmental concept to connect seemingly disparate syndromic phenotypes as outlined above and to potentially regard them as LPM diseases ^21^.

*rbm8a* perturbation in zebrafish has increasingly pleiotropic defects as described here and in previous work ^14^, in line with Rbm8a protein function in the ubiquitously deployed EJC that regulates diverse mRNA biology ^9,12,78,129^. Elegant genetic experiments have documented how the 3’ UTR intron-containing *foxo3b* mRNA that encodes a pro-apoptotic transcription factor is stabilized in *rbm8a*-mutant embryos, while double *rbm8a;foxo3b* mutants showed a significant rescue of the motor neuron defects ^14^. Not surprisingly, recessive null combinations of human RBM8A mutations have never been reported and are likely not viable; all genetically investigated TAR patients reported to date harbor at least one hypomorphic allele of *RBM8A*, suggesting reduced EJC levels or activity result in the combination of syndromic phenotypes ^1,5,6,130^. The combination of waning maternal contribution, morpholino-based dose attenuation, and our hypomorphic *rbm8a^Δ3^* allele in zebrafish provides *in vivo* means to approximate the assumed hypomorphic reduction of RBM8A levels in human TAR patients. In line with morpholino guidelines ^68^, *MO-rbm8aATG* injection morphologically phenocopy *rbm8a^Δ5/Δ5^*, reduces Rbm8a protein levels, is rescued with human RBM8A mRNA while expression lasts, and can be titrated for genetic interactions (**Fig. 1,2,6, Supplementary Fig. 1,2**).

Our observations that different PCP perturbations result in select hematopoietic defects (**Fig. 3,6**) provide a first mechanistic direction to consolidate the TAR defects. Our broad mRNA ISH panel for several developmental markers documented that *rbm8a*-mutant embryos do not feature broad LPM anomalies (**Supplementary Fig. 2**), and do not show the full complement of reported TAR phenotypes early in development ^2,5,6^. Notably, the typical limb and joint anomalies associated with the syndrome ^1,2^ cannot be fully modeled in zebrafish due to the evolutionary differences in extremity formation, limiting the use of zebrafish to model TAR Syndrome beyond basic developmental mechanisms. Nonetheless, our cell counting of *cd41:EGFP*-positive thrombocytes using two different methods including our new whole-larvae imaging (**Fig. 2, Supplementary Fig. 2**) provides evidence that *rbm8a* perturbation in zebrafish reduces overall thrombocyte numbers, while their ability to move, adhere and aggregate at sites of endothelial injury is not significantly perturbed (**Supplementary Fig. 2**). We further do not see evidence for disrupted endothelium mechanisms in our *rbm8a* mutants, as our ISH using *fli1a* shows no difference in endothelium/blood vessels (**Supplementary Fig. 2**).

Together, our data indicates that perturbed *rbm8a* affects the regulation of thrombocyte numbers and not their functionality, as appears to be the case for TAR syndrome patients ^2,5^. While TAR cases are rare and severe thrombocytopenia precludes standard platelet aggregometry, limited previous work has also found no overt functional defects in thrombocytes of TAR patients ^131^.

Convergence and extension in an embryo influences nearly all tissues and overall body architecture ^60,89,132^. Our morphometric analysis of 6-11 somite stage *rbm8a*-deficient embryos showed that *rbm8a^Δ5/Δ5^* mutant and *rbm8a* morphant embryos displayed phenotypes consistent with convergent and extension phenotypes (**Fig. 5**), even though the phenotypic characteristics were milder. The somites in classic convergent and extension mutants, such as *tri*/*vangl2*, amongst others, are generally wider; for example, in 7 somite stage *tri*/*vangl2* mutants, the somites are 1.27x wider than those in stage-matched wild type embryos ^90,133–136^. At the same timepoint, we observed that somites in *rbm8a^Δ5/Δ5^* mutants are 1.15x wider (**Supplementary Data 10**). Nonetheless, this paraxial mesoderm phenotype seems to be further amplified into the adjacent LPM (**Fig. 4**), which undergoes dramatic cellular and structural changes during convergence and extension movements, resulting in condensed bilateral stripes at the periphery of the embryo. The subsequent midline migration of hematopoietic, endothelial, and kidney progenitors is sensitive to perturbations of guidance cues, yet the exact mechanism(s) that trigger and drive midline migration remain incompletely resolved. PCP signaling defects prevent condensation of cells across all germ layers along the anterior-posterior axis, keeping especially the LPM as lateral-most cells widely dispersed ^55,60,89,91,137,138^.

How could attenuation of Wnt/PCP signaling influence hematopoietic progenitors? One possible explanation could be that as hematopoietic progenitors migrate as medial-most LPM stripe and are exposed to different signals and cellular interactions during their migration ^20^, attenuated PCP could interrupt hematopoietic progenitor arrangement and migration speed. This model is in line with our observations of altered *scl:EGFP*-expressing LPM in *rbm8a* and *vangl2* perturbations (**Fig. 4**). However, in both *rbm8a* and *vangl2* mutants hematopoietic progenitors do still form, albeit with altered expression of individual regulatory genes (**Fig. 7**). During their medial migration, LPM cells are in constant contact with endoderm and overlaying somites that have been implicated in influencing hematopoietic progenitor patterning and fate potential ^62,63,139,140^. Non-canonical Wnt signaling has been implicated in controlling somite-expressed Notch ligands that are required for hematopoietic stem cell progenitor emergence ^62^. While we do see a reduction of *runx1*-positive progenitors (**Fig. 7**), they are perturbed already before *runx1* expression refines to HSPCs in the dorsal aorta after 24-26 hpf, suggesting an earlier influence on hematopoietic progenitors following attenuated Wnt/PCP in *rbm8a*-deficient embryos. Nonetheless, the involvement of non-canonical Wnt/PCP signaling at different steps of hematopoiesis might contribute to the selective thrombocyte anomalies observed in TAR patients, warranting further experimental follow-up.

We therefore hypothesize that, following Rbm8a perturbations, attenuated PCP through retention of mis-spliced mRNAs encoding non-canonical Wnt/PCP components affects migration or relative position of hematopoietic progenitors within the embryo as they migrate through signaling gradients. Notably, mice with defects in the EJC component Srsf3 also display thrombocytopenia ^141^. The progressive recovery of the thrombocytopenia phenotype in TAR patients is compatible with an early defect in embryonic or fetal hematopoiesis when transient or intermediate progenitor waves are forming; the high plasticity and clonal expansion even of potentially reduced hematopoietic stem cells could subsequently replenish the diminished megakaryocyte levels during definitive hematopoiesis ^1,2,25,142–145^. The reduced *gfi1aa* expression (**Fig. 7**) and later reduced numbers of *cd41:EGFP*-expressing thrombocytes in our quantifications following *rbm8a* and PCP perturbations show signs of compensation over time (**Fig. 2,6**), providing an experimental starting point to pursue this model.

Connecting morphological alterations by attenuating non-canonical Wnt/PCP signaling as major pathway involved in embryo morphogenesis provides a possible mechanistic link to the impact on LPM-derived structures in TAR Syndrome and possibly other congenital conditions. While we here focus on consequences of perturbed *rbm8a* on hematopoiesis, non-canonical Wnt/PCP disruptions have the potential to affect a variety of tissues ^49^. PCP signaling does not elicit an obvious transcriptional response, confining effects of its perturbation to morphological defects; however, we hypothesize that incorrect or delayed migration or altered arrangement of LPM progenitors could manifest as defects in individual downstream fates by altering the spatio-temporal dynamics of exposure to developmental signals and cell-cell interactions. Such a model would be consistent with recent observations linking emerging cell fates to overall morphogenesis that is a prerequisite for the correct formation of patterning gradients ^146^. In addition to zebrafish not forming a radius bone, future work is warranted to decode the basic impact on heart, forelimbs (pectoral fin), kidney, and other LPM-derived structures upon *rbm8a* perturbation. PCP defects have been linked to anomalies in each of these structures in mice ^49,52,54,64,103,147–150^, further suggesting defective PCP signaling in LPM patterning as corroborating developmental link for the TAR syndrome phenotypes.

## MATERIALS AND METHODS

All research described herein complies with all relevant ethical regulations as reviewed and approved by the University of Colorado Anschutz Medical Campus, the University of Zurich, the MDC Berlin, and Baylor College of Medicine.

### Zebrafish husbandry

Zebrafish (*Danio rerio*) husbandry and experiments were performed according to IACUC regulations at the University of Colorado Anschutz Medical Campus (protocol number 00979) and the University of Michigan (protocol number PRO00010679), FELASA guidelines ^151^, the guidelines of the Max-Delbrück Center for Molecular Medicine in the Helmholtz association, and the local authority for animal protection (Landesamt für Gesundheit und Soziales, Berlin, Germany). The ‘Principles of Laboratory Animal Care’ (NIH publication no. 86– 23, revised 1985) as well as the current version of German Law on the Protection of Animals and EU directive 2010/63/EU on the protection of animals were followed. All zebrafish were raised, kept, and handled according to established protocols if not noted otherwise ^152^.

### Transgenic zebrafish lines

The following established transgenic lines were used in this study: *Tg(scl:EGFP)^zf255^* ^82^, *Tg(cd41:EGFP)* ^69^, and *Tg(RGB)* ^153^ (combined transgene insertions from *Tg(-6.35drl:EGFP)^cz3331^* ^102^, *Tg(lmo2:loxP-dsRED2-loxP-EGFP)^rj2^* ^154^, and *Tg(myl7-loxP-AmCyan-loxP_ZsVenus)^fb2^* ^155^.

### sgRNA template generation, RNP complex assembly and injection

Oligos were obtained from Sigma/Millipore and Life Technologies as standard primers except where otherwise noted. A short guide RNA (sgRNA) template was generated by oligo-based PCR amplification (Bassett et al., 2013) using the sgRNA forward primer containing a *T7* promoter sequence *5’-GAAATTAATACGACTCACTATAGGGAGGCGAAGACTTTCCTAGTTTTAGAGCTAGAAATAGC-3’* and the invariant reverse primer *5’-AAAAGCACCGACTCGGTGCCACTTTTTCAAGTTGATAACGGACTAGCCTTATTTTAACTTGCTATTTCTAGCTCTAAAAC-3’* (PAGE-purified). A primer extension reaction was performed using Phusion DNA polymerase (M0530L, NEB) followed by QIAquick purification (28106, Qiagen) with elution of the PCR product in DEPC-treated water. *In vitro* transcription (IVT) of the sgRNA-based PCR template was performed using the MAXIscript T7 Kit (AM132, Ambion) with an overnight reaction run at 37°C, followed by ammonium acetate precipitation or clean up via RNA isolation kits as per the manufacturer’s instructions and as described previously ^156,157^. The only change to the manufacturer’s protocol of the MAXIscript T7 Kit was to add 100mM NTPs instead of the recommended 10 mM as this greatly increases sgRNA yield. Before use, the sgRNA was quality controlled on a denaturing 2.5% MOPS gel. The CRISPR-Cas9 ribonucleoprotein complex (RNPs) was assembled using appropriate salt concentrations to keep the RNPs solubilized (Burger et al., 2016). Salt-solubilized RNPs were then injected at sub-optimal concentrations to increase viability and founder identification (Burger et al., 2016).

### Mutant zebrafish lines

*rbm8a* mutant alleles were generated as per our previous work ^158^ using the target site 5’-*GGGAGGCGAAGACTTTCCTA-5’* as proposed by CHOPCHOP ^159^. Details about sgRNA template generation and RNP complex assembly and injection are described in the Supplemental Data. The two alleles *rbm8a^Δ5^* (also described in Gangras et al., 2020) and *rbm8a^Δ3^* were selected. Both alleles were genotyped by sequencing PCR products using primer *rbm8a fwd 5’-AAACAGCAGACGGCGAGG-3’* and primer *rbm8a rev 5’-GCTGAATCACTACAACGCG-3*’. Additionally, the *rbm8a* wild type allele was genotyped by restriction digest of the PCR product with Xmn1 (R0194S, NEB) and the *rbm8a^Δ3^* allele by Hinf1 (R0155S, NEB) while the *rbm8a^Δ5^* allele stayed intact upon Xmn1 or Hinf1 digest.

Further, the established *vangl2^vu67^* mutant was used^88^ (a kind gift from Dr. Liliana Solnica-Krezel). Embryos were genotyped by sequencing PCR products using primer *vangl2 fwd 5’-ATTCCCTGGAGCCCTGCGGGAC-3’* and primer *vangl2 rev 5’-AGCGCGTCCACCAGCGACACAGC-3*’ or restriction digest of the PCR products with Alu1 (R0137S, NEB). The *vangl2* wild type allele stayed intact while the *vangl2^vu67^* mutant allele was identified by a digested PCR product.

### Morpholino injections and rescue experiments with capped mRNA

Capped mRNA for mutant rescue was generated using the mMESSAGE mMACHINE™ SP6 Transcription Kit (AM1340, Ambion) as per the manufacturer’s instructions. As templates, *pCS2+:EGFP*^160^ and *pCS2FA:hs_RBM8A* were used. To generate the original *pCS2FA:hs_RBM8A*, the human *RBM8A* ORF was amplified from HEK293T cDNA by PCR with primer *fwd 5’-TTA<u>GGATCC</u>ATGGCGGACGTGCTAGATC-3’* containing a *BamHI* site (underlined) and primer *rev 5’-TT<u>GCTAGC</u>TCAGCGACGTCTCCGGTCT-3’* containing a *NheI* site (underlined). The PCR product was cloned with BamHI (R0136S, NEB), NheI (R3131S, NEB) restriction digest into Tol2kit vector *396* (*pCS2FA:Tol2*)^161^ cut with BamHI and XbaI (R0145S, NEB).

Morpholinos were obtained from Gene Tools (Philomath, USA) and prepared as a 1 mM stock solution according to the manufacturer’s instructions. To target *rbm8a*, a translation blocking morpholino, *MO-rbm8aATG* with sequence *5’-GAAGATCCAGTACGTCCGC<u>CAT</u>GA-3’* (sequence complementary to predicted start codon is underlined), was designed and diluted as indicated. To achieve full *rbm8a* knockdown a dilution of 1:100 was used, and is referred to as optimal dilution. To achieve suboptimal rbm8a knockdown, dilutions of 1:125, 1:150, 1:175, 1:200, and 1:500 were used. Of note, optimal and suboptimal dilutions differed between labs, and are indicated in the respective figures. All other morpholinos used in this study have been published previously and morpholino details and injection procedures are described below.

Rescue experiments were performed by injecting *MO-rbm8aATG* (1:100 dilution) and 100 ng/µL capped *hs_RBM8A(altered)* mRNA and *EGFP* mRNA as control. As the codons encoding the N-terminus of zebrafish and human RBM8A protein are identical in the respective mRNAs and therefore also human *RBM8A* mRNA is recognized by the *MO-rbm8aATG*, 5’ codons in the morpholino-targeted region were altered to synonymous codons where possible: wild type human *RBM8A 5’-<u>ATG</u>GCGGACGTGCTAGATCTTCACGAGGCTGGG-3’* to *5’-<u>atg</u>gcAgaTgtCTtGgaCTtGCAC-3’* (alterations in capital letters, resulting in synonymous codons, *<u>ATG</u>* start codon underlined in both sequences). The resulting *RBM8A(altered)* ORF was ordered from Twist Biosciences and swapped into *pCS2FA:hs_RBM8A* with flanking BamHI and NheI sites as above, resulting in *pCS2FA:hs_RBM8A(altered).* mRNA was generated and injected as outlined above.

The embryos were observed for phenotypes at 24, 48 and 72 hpf and representative images were taken with a 10x lens on a Zeiss LSM880. Observed phenotypes were plotted in percent using GraphPad Prism (10.0.3).

### Additional morpholinos and injection procedures

*MO1-vangl2* (ZFIN ID: *ZDB-MRPHLNO-041217-5*, 5’-*GTACTGCGACTCGTTATCCATGTC*-3’), *MO2-wnt5b* (ZFIN ID: *ZDB-MRPHLNO-051207-1*, *5’-TGTTTATTTCCTCACCATTCTTCCG-3’*), *MO2-wnt11f2* (formerly known as *MO2-wnt11* or *wnt11-MO*, ZFIN ID: *ZDB-MRPHLNO-050318-3*, 5’-*GTTCCTGTATTCTGTCATGTCGCTC*-3’) and *MO1-fzd7a* (ZFIN ID: ZDB-*MRPHLNO-050923-5*, *5’-ATAAACCAACAAAAACCTCCTCGTC-3’*) which we have described previously and validated^52,68,162–165^, have been used at indicated dilution. Lastly, the Gene Tools pre-made standard control oligo morpholino (100 nmol; diluted to 1:200) was used.

Prior to injections, the 1mM morpholino stock solution was heated up to 65 °C, vortexed for 5 minutes and diluted to achieve the desired concentrations. The diluted morpholino solution was again heated to 65 °C and vortexed for 10 minutes before injecting. All morpholino solutions were stored at room temperature.

Microinjections were performed using MPPI-3 microinjectors (Applied Scientific Instrumentation) together with a Narishige micromanipulator (MN-153). Morpholino oligonucleotides were injected into the yolk of one-cell stage zebrafish embryos. The injection volume was calibrated to the final volume of 1 nL in total. In case of double knockdown approaches to study genetic interactions, both morpholinos were injected sequentially. Embryos were kept in E3 embryo medium under standard laboratory conditions at 28.5 °C.

### Zebrafish protein sample collection

Zebrafish embryos were collected at indicated developmental stages (tailbud stage, 10 somite stage, 16 somite stage, 24 hpf). Individual embryos were transferred into tubes kept on ice filled with 200 µL cell dissociation solution (CDS; 5 ml of Ringers’ solution (96724, Sigma), 1 tablet of Complete Mini EDTA-free (11836153001, Roche). The embryos in CDS were flash frozen in liquid nitrogen and samples were stored at -80 °C until they were genotyped and could be combined according to their genotype for further usage. For genotyping, a few cells from individual embryos were collected and genotyped as described above.

After combining 10 embryos with the same genotype for each developmental stage, the samples were centrifuged at 1500 rpm for 2 minutes at 4 °C. The tubes were turned by 180° and centrifuged for 2 additional minutes. The supernatant was removed, 200 µL CDS was added, and centrifuged again as above. Afterwards, the supernatant was again removed, and the samples centrifuged once more with additional removal of the remaining supernatant using fine gel loading tips. Then, 20 µL 2x SDS Laemmli buffer (1610747, BioRad) supplemented with 5% 2-betamercaptaethanol (M3148, Sigma) was added to each sample, vortexed rigorously, and cooked for 5 min at 95 °C.

### Electrophoresis and immunoblot analysis

Samples were loaded on a precast Mini-PROTEAN TGX gel (4569034, Biorad) and electrophoresis was performed as per the manufacturer’s instructions. Afterwards, the gel was transferred onto a PVDF membrane (1704272, Biorad) using the trans-blot turbo transfer system (1704150, Biorad).

Zebrafish Rbm8a and Tubulin proteins were detected using standard immunodetection protocols, in short: blocking with 1XTTBS 5% BSA for 1 hour, overnight incubation with anti-Rbm8a (1:1000; 05-1511, Millipore), anti-Tubulin (1:1000; 05-829, Millipore) in 1XTTBS 5% BSA at 4 °C, followed by 3 x 5 min wash in TTBS and 30 min incubation with IRDye 800CW goat anti-Mouse (1:10000; 926-32210, Li-Cor) in 1XTTBS 5% BSA at room temperature followed by 3 x 5 min wash in 1XTTBS. Images of Western blots were obtained using an Odyssey CLx imager (Li-Cor) and quantified using ImageJ (https://www.yorku.ca/yisheng/Internal/Protocols/ImageJ.pdf). All full gel images are shown **Supplementary Fig. 1**.

### *cd41:EGFP*-positive cell counting

For live imaging of 3 and 6 dpf *cd41:EGFP* zebrafish larvae, the heartbeat was stopped with 30 mM 2,3-butanedione monoxime (BDM; B0753, Sigma) in E3, in addition to anesthesia with Tricaine. Images of the zebrafish larvae used for thrombocyte counting were taken on a Leica M205 FA with Plan Apo 1.0x lens. 3 dpf larvae were imaged directly in E3+BDM+Tricaine on a glass plate, while the 6 dpf larvae were placed on a petri dish covered with a 2% agarose layer. Larvae were placed anterior to the left. Images were taken consistently at 2x zoom for groups of two-four zebrafish larvae at 3 dpf and 3x zoom for a single zebrafish larvae at 6 dpf. For analysis, the images were exported from LAS Leica, and using Fiji/ImageJ macro ^166^, the GFP channels were separated. At 3 dpf, the cells were counted manually by adjusting brightness and contrast as needed, using the Fiji/ImageJ cell counter plugin (https://imagej.nih.gov/ij/plugins/cell-counter.html). At 6 dpf, a second Fiji/ImageJ macro was run that thresholds and automatically counted the cells. Representative images of 6 dpf larvae for figure panels were taken on a Zeiss LSM880 with 10x lens. In 6 dpf larvae, Fiji/ImageJ counts were manually recounted for verification. Individual *cd41:EGFP* values, means, and ± S.D. were plotted using GraphPad Prism (10.0.3).

Macros to run on imaging files in an individual folder:

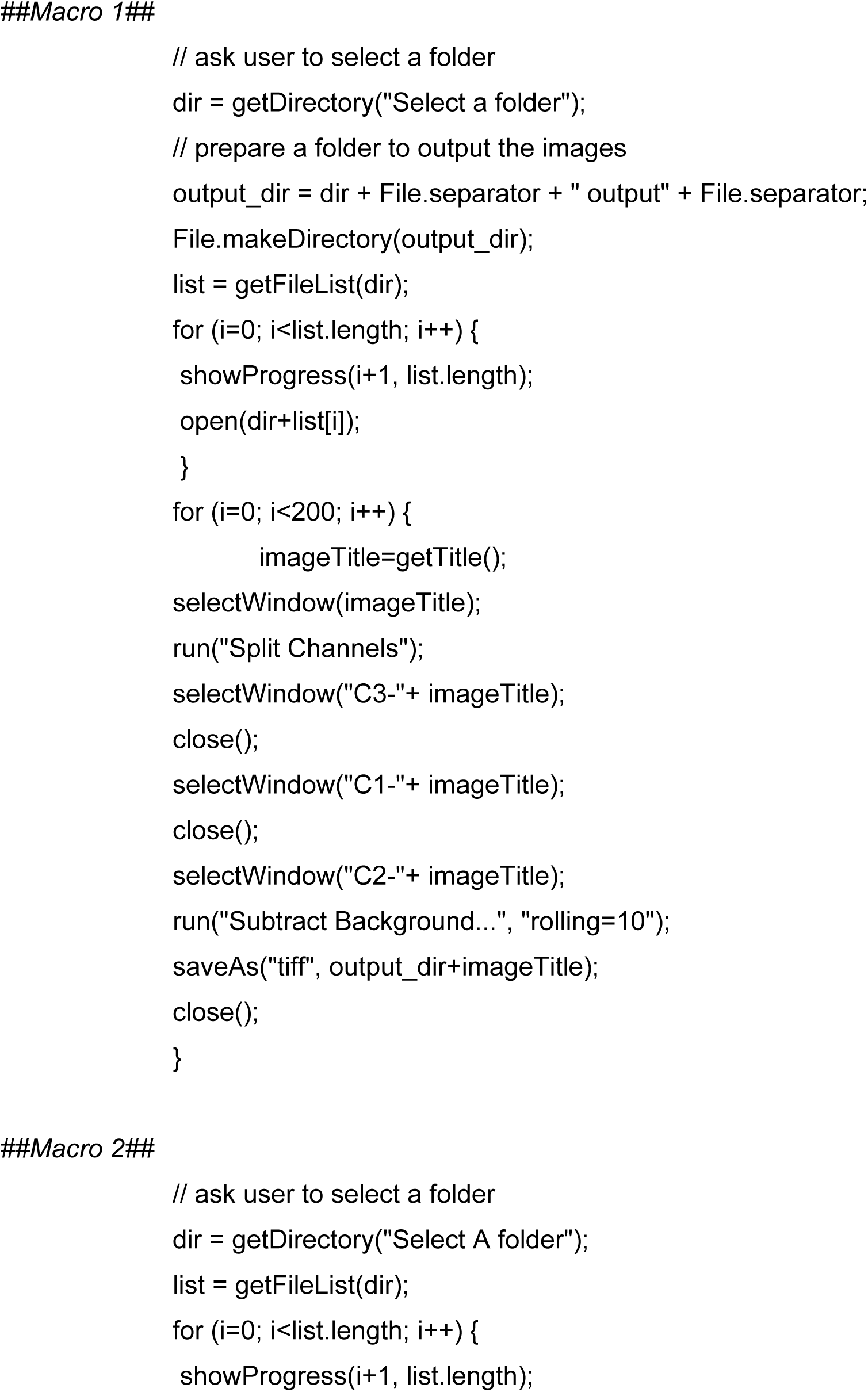
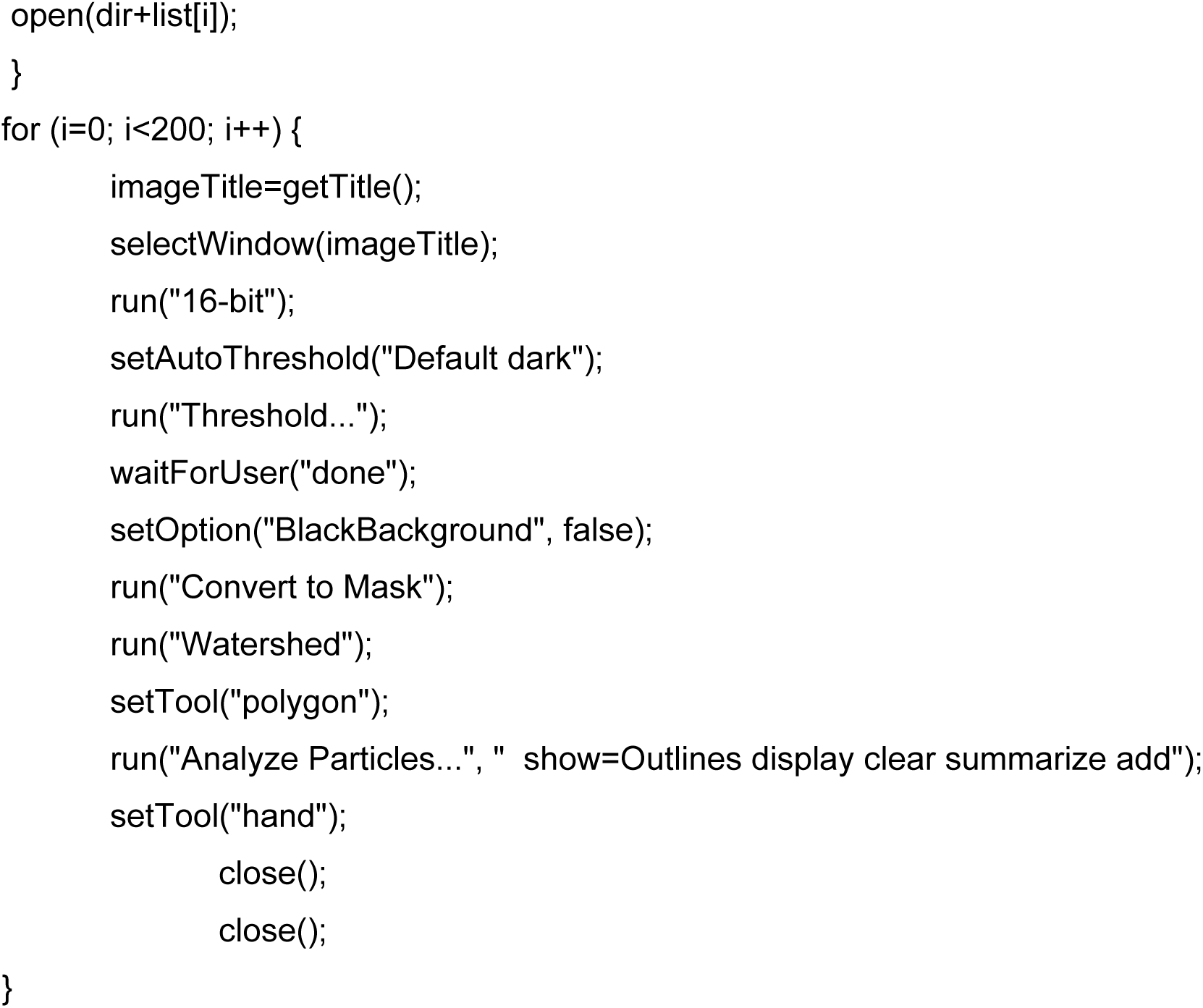

Upon completion, toggle the top bar until all *cd41*:EGFP-bright cells are covered, press “ok” in dialog window. Resulting data was processed in Excel for next steps.

### Circulating thrombocyte video quantitation and functional analysis

Circulating *cd41:EGFP-*positive thrombocytes were measured as previously described^73,74^. 6 dpf larvae *cd41:EGFP* were anaesthetized in tricaine and submersed in a solution of 0.7% low-melt agarose. 4 to 6 larvae were aspirated into 1.5-1.8 mm diameter capillary tubes (Pyrex) using a manual pipette pump, each separated by an air bubble. The capillary was mounted on modeling clay, submerged in 1x E3 medium in a 10 cm Petri dish, and then placed under a stereomicroscope (Leica) with an Olympus DP22 2.8-megapixel digital camera. Magnification was set at 85x and a lateral view of the anterior tip of the larva to the end of yolk sac extension was visualized. Fluorescent videos were captured for a duration of one minute and then analyzed using VirtualDub (v1.10.4) and Fiji ^166^. The first 1000 frames were quantified, averaged, and then background subtracted to eliminate non-circulating cells. Finally, zebrafish larvae were removed from the capillary, lysed, and genotyped.

Laser-mediated arterial endothelial injury was performed using a pulsed-dye laser system (Micropoint, Andor Technology), as previously described^74,75^. 6 dpf zebrafish larvae carrying the *cd41:EGFP* transgene were anesthetized in tricaine, positioned laterally in 0.8% low-melt agarose on glass cover slips and visualized on an inverted microscope (Olympus 1X73) using a 20x objective. The endothelium of the dorsal aorta was ablated at the fifth somite posterior to the anal pore. The time to occlusion was recorded up to 120 seconds and the number of *cd41:EGFP-*positive thrombocytes adhering at the site of injury during that period was recorded. After laser injury, zebrafish larvae were recovered from the agarose, lysed, and genotyped. Individual values, means, and ± S.D. were plotted using GraphPad Prism (10.0.3).

### Bulk RNA sequencing data analysis

RNA sequencing reads were initially quality checked with FastQC (https://www.bioinformatics.babraham.ac.uk/projects/fastqc/) and MultiQC^167^ and deemed of high quality. We quantified transcript-level expression using Salmon (version 0.8.2) ^168^ against the Ensembl GRCz10.87 catalog ^169^ and performed gene-level differential expression using tximport (version 1.4.0) ^170^ and edgeR (version 3.18.1) ^171^ using the quasi-likelihood F test ^172^ and a Benjamini-Hochberg multiple testing correction ^173^; thresholds were set at FDR <= 0.05. Volcano plots were produced using the ggplot2 (version 2.2.1) package. RNA-seq reads were also mapped against the GRCz10 reference genome with STAR (version 2.5.1b_modified) ^174^, using the Ensembl GRCz10.87 catalog. Coverage tracks (in bigWig format) were created with bedtools (version 2.17.0)^175^ and bedGraphToBigWig (version 4)^176^ and plotted with Gviz (version 1.20.0) ^177^. Differential exon/intron usage analysis was performed using DEXSeq (version 1.22.0) ^79^. Transcripts contributing less than 5% of the abundance for a given gene in all samples were removed from the gtf file ^178^ and a gff file with disjoint exon bins were generated from the remaining transcripts using DEXSeq. Finally, introns (regions of a gene locus not belonging to any annotated transcript) were explicitly added as separate bins in the gff file before quantification and differential exon/intron usage analysis with DEXSeq. To explore the RNA-seq data interactively, we created the R/Shiny-based RNAseq Explorer app (link: http://imlspenticton.uzh.ch:3838/mosimann_p2452/). The app contains tabs for various visualizations and analyses: individual gene expression levels (by typing the gene name), a PCA plot for an overall view of the similarity of gene expression profiles, normalized base-level coverage tracks for individual genes, differential expression results, differential transcript usage results, and gene set enrichment analyses.

### Light sheet sample preparation, microscopy, and image analysis

The Zeiss Z.1 microscope equipped with a Zeiss W Plan-Apochromat 20×/0.5 NA objective was used for all light sheet microscopy. Embryos within the chorion were embedded in 1% LMA in a 50 µL glass capillary. For the multi-angle imaging data sets, the four individual angles per embryo were manually registered before applying the Fiji Multiview Reconstruction and BigStitcher plugins ^166,179^ for fusion and deconvolution of the images. Images and movies were further processed using Fiji/ImageJ and Imaris (9.7)^180^.

Morpholino-injected embryos were raised at 28 °C and imaged in the Z.1 from 4 somite stage to 12 somite stage, alongside uninjected *scl*:EGFP controls. In Imaris, the baseline and background were subtracted and surface rendering was performed to automate volume and area determination of the fluorescent signal.

Quantifications of area and volume from wild type, *MO-rbm8aATG* and *MO1-vangl2*-injected embryos were averaged for each timepoint. Average values and ± S.D. were plotted using GraphPad Prism (10.0.3).

### Whole mount *in situ* hybridization

Whole-mount in situ hybridization (WISH) was performed according to the Thisse lab protocol ^181^. Zebrafish embryos were fixed overnight in 4% PFA (47608, Sigma) in PBS (EC-A500, Life Sciences), dechorionated, rinsed in PBS and dehydrated into methanol. Samples were then rehydrated into PBT (PBS + Tween (P1379, Sigma), treated with Proteinase K (1:10000) (P8108S, NEB) for 2 min, washed in PBT and postfixed in 4% PFA/ 0.2% glutaraldehyde (G5882, Sigma) in PBS for 20 min at RT. Then the samples were incubated for at least two hours in hybridization solution with 50% formamide (in 0.75 M sodium chloride, 75 mM sodium citrate, 0.1% Tween-20 (P1379, Sigma), 50 mg/mL heparin (H3393, Sigma), and 200 mg/mL tRNA (AM7119, Invitrogen)) at 70°C, then hybridized overnight at 70°C with antisense probes diluted approximately 1 ng/ml in hybridization solution. Samples were washed gradually into 2X SSC buffer (0.3 M sodium chloride, 30 mM sodium citrate or AM9763, Invitrogen), and then gradually from SSC to PBT. Samples were blocked at room temperature for several hours in PBT with 2% goat or sheep serum (S2263, Sigma) and 2 mg/mL bovine serum albumin (BSA, 196941, Fisher), then incubated overnight at 4°C with anti-DIG antibody (11093274910, Roche) at 1:5000 in blocking buffer. Samples were rinsed extensively in PBT and staining buffer (100 mM Tris-HCl pH 9.5, 50 mM MgCl2, 100 mM NaCl, 0.1% Tween 20, 0.3mg/mL tetramizole hydrochloride (T1512, Sigma) in dH20), prior to staining with BM Purple AP staining solution (62321800, Roche). Staining was stopped by washing embryos in PBT.

### Antisense riboprobes for *in situ* hybridization

The templates for all *in situ* hybridization (ISH) probes were generated by PCR amplification using primers and complementary DNA (cDNA) or linearized plasmids. First-strand cDNA was generated from pooled zebrafish RNA isolated from different developmental stages using SuperScript III first-strand synthesis kit (18080051, Invitrogen). Subsequently, DNA templates for ISH probes were generated using first-strand cDNA as a PCR template and primers as specified for each gene of interest; for *in vitro* transcription (IVT), either the *T7* (*5′-TAATACGACTCACTATAG*-3′) or *T3* (5′-*AATTAACCCTCACTAAAG*-3′) promoter sequence was added to the *5′* ends of the reverse primers. PCR reactions were performed under standard conditions as per manufacturer’s protocol using Phusion DNA polymerase.

The following primer or plasmids to generate templates for ISH probes were used: *kdrl* (using *pBS-kdrl* (a kind gift from Dr. Leonard I. Zon), linearized with EcoRI (R0101S, NEB), T7 polymerase for IVT)^182^; *sox7* (using primer *fwd 5’-CGACCAAAACTCCCTTCCG-3’* and primer rev *5’-AAAA<u>TAATACGACTCACTATAG</u>GGTTGTTGTAGTAGGCTGC-3’*, T7 polymerase for IVT), *gata1* (using primer f*wd 5’-TGGGAAAGACAGTCCCAGG-3’* and primer *rev 5’-AAAAA<u>TAATACGACTCACTATAG</u>GGCCTTCACACTAGTGTGGG-3’*, T7 polymerase for IVT); *runx1* (using *pBS-runx1a*, linearized with HindIII (R0104S, NEB), T7 polymerase for IVT)^183^ (a kind gift from Dr. Teresa V. Bowman); *gfi1aa* (using primer *fwd 5’-TTATCATCAGCCCCGTTACC-3’* and primer *rev 5’-AAAA<u>TAATACGACTCACTATAG</u>GGAATGGACGGCTTTATGTTGC-3’*, T7 polymerase for IVT)^42^; *gfi1b* (using primer *fwd 5’-ACCAACCTCAAACGAGAGC-3’* and primer *rev 5’-AAAA<u>TAATACGACTCACTATAG</u>GGATTGTCCATCAACTTCTGTC-3’*, T7 polymerase for IVT) ^42^, *myoD* (linearized with BamHI, T7 polymerase for IVT) ^99^, *dlx3b* (in pBS, linearized with SalI, T7 polymerase for IVT) ^100^, *tbxta* (in pBS-KS, linearized with XhoI, T7 polymerase for IVT) ^97^, and *hgg1* (linearized with EcoRI, T7 polymerase for IVT) ^101^.

Antisense ISH probes were transcribed from their template overnight at 37 °C using either MAXIscript T7 Kit (AM132, Ambion) or MAXIscript T3 Kit (AM1316, Ambion) and digoxigenin (DIG)-labeled NTPs (11277073910, Roche) as per the manufacturer’s instructions. The resulting RNA probe was treated with TURBO DNase (AM2238, Invitrogen), precipitated with lithium chloride (9480G, Invitrogen) and EtOH, dried, rehydrated in 10-20 µL DEPC-treated water and stored at -80 °C until further use.

### DNA extraction from fixed WISH zebrafish embryos for genotyping

Residual PBT was removed from embryos and replaced with 50 µL 300 mM NaCl. The sample was incubated for 4 hours at 65 °C for reverse crosslinking. Afterwards, NaCl was removed and replenished with 50 µl ELB (50 ml: 500 µL 1M Tris pH 8.0, 2.5 ml 1M KCl, 150 µL Tween-20 (P1379, Sigma), 47 ml water) and 4 µL 10mg/ml Proteinase K (P8108S, NEB). The samples were incubated for 8 hours at 55 °C followed by 10 min at 98 °C to inactivate the Proteinase K. The samples were centrifuged and stored at -20 °C until genotyping procedure (see above).

### Image analysis for morphometric measurements

WISH-stained embryos with *myoD*, *dlx3b*, *tbxta*, and *hgg1* were transferred into a clear flat-bottom dish with methylcellulose and imaged using a Nikon SMZ18 stereomicroscope. After imaging, embryos were transferred back in PBT for storage at 4 °C or immediately genotyped as described above.

ImageJ was used to visualize and manipulate images of WISH-stained embryos and measurements were made manually using the line and angle tools ^166,184^. Axis length and angle were collected by measuring from the anterior to posterior aspects of the *dlx3b* expression domain in lateral-view images. Neural plate width was collected by measuring from one side of the *dlx3b* expression domain to the other side at the level of the otic placodes and beginning of the notochord in dorsal-view images. Somite width was taken from dorsal-view images by measuring the left to the right side of the *myoD* expression domain at the same last three somite pairs per embryo.

Quantification of axis length, axis angle, neural plate width, and somite width across conditions were averaged for each timepoint. Average values and ± S.D. were plotted using GraphPad Prism (10.0.3).

### Image analysis of hematopoietic markers and quantification of *ISH* signal

WISH-stained embryos with *kdrl*, *sox7*, *gata1*, *runx1*, *gfi1aa*, and *gfi1b* were transferred into a clear flat-bottom dish with glycerol and imaged using a Leica M205 dissecting scope with TL3000 Ergo base, swan neck lamps, Planapo 1.0x M-series objective, and FLEXACAM C1 camera.

Quantification of the ISH signal for *gata1, gfi1aa*, and *gfi1b* was done as previously described using ImageJ ^185^. Briefly, after imaging the WISH-stained embryos with consistent parameters, the images were inverted to negative and converted to 8-bit grayscale. A region of interest (ROI) containing the ISH expression signal was drawn manually for each embryo using the freehand selection tool in ImageJ. To define the background, a second ROI with the same shape and area was created in an unstained region of the trunk, above the ROI with the signal. A value for each region was determined by measuring the average pixel intensity. The pixel intensity of the ISH signal was determined by subtracting the value of the background region from the value of the stained region. Individual data points, means, and ± S.D. were plotted using GraphPad Prism (10.0.3). Quantification of the ISH signal intensity in AGM, PBI, and neuron territories in *runx1*-stained embryos was done by giving the signal intensity a score from 1 to 5 by three independent researchers, totaling four counts per embryo. Counts were averaged and are presented in the graph as individual data points. Individual data points, means and ± S.D. were plotted using GraphPad Prism (10.0.3).

### Experimental study design

All assays were treated with identical experimental conditions across species and performed at least twice or more times. All attempts at replication were successful.

No data were excluded in the zebrafish studies.

Data analyses of the *cd41*-transgenic reporter quantification was based on morpholino injections into zebrafish embryos and based on defined genotypes of zebrafish embryos from crosses. No other randomizations were applicable.

Data collection for the *cd41:EGFP*-positive cell counting was blinded to avoid researcher bias and counted by at least two independent researchers. Data collection for the circulating thrombocyte video quantitation and functional analysis was recorded by a researcher blinded to the genotype. Data collection of ISH analyses were blinded and morphometrics and ISH signal intensity were measured by at least two independent researchers.

Zebrafish embryos were not selected by gender as sex determination happens later in development.

### Statistical analysis

The authors declare that key measures of statistics and reproducibility are built into the work throughout. For the zebrafish assays, sufficient embryos were analyzed to achieve statistical significance based on previous experience in these studies. Experimental sample sizes were chosen by common standards in the field and in accordance with solid phenotype designation.

### Data and code availability

All mRNA sequencing data has been deposited at ArrayExpress with accession E-MTAB-12503. The related GitHub repository (link: https://github.com/markrobinsonuzh/rbm8a-rnaseq-explorer) hosts code and R objects to support the R/Shiny app for exploring *rbm8a* mutant RNA-seq experiments interactively. All primary data from the *cd41:EGFP*-positive cell counts, SPIM imaging of the *scl:GFP*-positive territory to determine area and volume, ISH morphometrics and quantifications are included as tables in the Supplementary Data. The western blots are included in Supplementary Fig. 1.

The code for the R/Shiny-based RNAseq Explorer app and instructions for its use are available at https://github.com/markrobinsonuzh/rbm8a-rnaseq-explorer, with track data downloadable from https://doi.org/10.5281/zenodo.7680271.

## ACKNOWLEDGEMENTS

We thank Christine Archer, Molly Waters, Nikki Tsuji, Ainsley Gilbart, and Olivia Gomez (CU Anschutz), as well as Vesna Barros, Lukas Obernosterer, and Yorgos Panayotu (UZH), and the aquatic facility staff at MDC for expert zebrafish husbandry support, Sibylle Burger and Cassie L. Kemmler for excellent technical assistance, Dr. Robert L. Lalonde for genotyping troubleshooting, Hannah R. Moran for support with quantifying *runx1 in situ* data, and past and current members of the Mosimann and Burger labs for critical discussions. We are grateful to the following colleagues for kind reagent sharing and input: Dr. Lilianna Solnica-Krezel for sharing *vangl2^vu67^* zebrafish; Dr. Teresa V. Bowman and Dr. Leonard I Zon for sharing the *pBS-kdrl* and *pBS-runx1* plasmids; Dr. Martin Gering for critical input on performing *gfi1aa* and *gfi1b in situ* hybridization; Dr. Martin Jinek for input on EJC interactions and the nature of the isolated *rbm8a* alleles; Dr. Guramrit Singh for pointers concerning functional Rbm8a antibody alternatives. Schematics that include Biorender.com elements are indicated in the corresponding figure legends.

This work has been supported by NIH/NIDDK grant 1R01DK129350-01A1 and an UZH URPP “Translational Cancer Research” seed grant to A.B.; a project grant from the SwissBridge Foundation and Additional Ventures SVRF2021-1048003 to C.M. and A.B.; a Swiss National Science Foundation (SNSF) professorship (PP00P3_139093) and Sinergia Grant (CRSII5_180345), a Marie Curie Career Integration Grant from the European Commission (CIG PCIG14-GA-2013-631984), the Canton of Zürich, the UZH Foundation for Research in Science and the Humanities, the UZH Science Alumni, Blutspende Zürich, National Science Foundation Grant 2203311, the University of Colorado School of Medicine, and the Children’s Hospital Colorado Foundation to C.M.; C.M. holds the Helen and Arthur Johnson Chair for the Cardiac Research Director; NIH/NIGMS GDDR T32 1T32GM141742 support to H.H.W.; SNSF grants 310030_175841 and CRSII5_177208 and University of Zurich’s Research Priority Program Evolution in Action to M.D.R..; the Helmholtz Association (ERC-RA-0047), DZHK and BMBF (German Center for Cardiovascular Research and German Ministry of Education and Research, respectively, project 81X3100110) to D.P.; an Additional Ventures SVRF grant to C.M., A.B., and D.P.; NIH/NIGMS grant R00HD091386 to M.L.K.W.; a National Hemophilia Foundation Judith Graham Pool Postdoctoral Fellowship Award to A.R.; NIH/NHLBI R35HL150784 to J.A.S.; J.A.S. is the Henry and Mala Dorfman Family Professor of Pediatric Hematology/Oncology.

## AUTHOR CONTRIBUTIONS

A.K., E.C., H.H.W., M.S.H., S.T.J., C.M., A.B., zebrafish experiment design and execution, manuscript writing; C.S., M.D.R., bulk RNA sequencing, data analysis, manuscript writing; J.S.M., M.L.K.W. convergence and extension morphometric measurements, data analysis, manuscript writing; K.M.M.A., D.P., PCP and genetic interaction experiment design and execution, data analysis, manuscript writing; A.R, J.A.S. video-based thrombocyte measurements and laser injury assay, data analysis and interpretation.

## COMPETING INTERESTS STATEMENT

J.A.S. has been a consultant for Sanofi, Takeda, Genentech, CSL Behring, and HEMA Biologics.

---

**Supplementary Figure 1:**
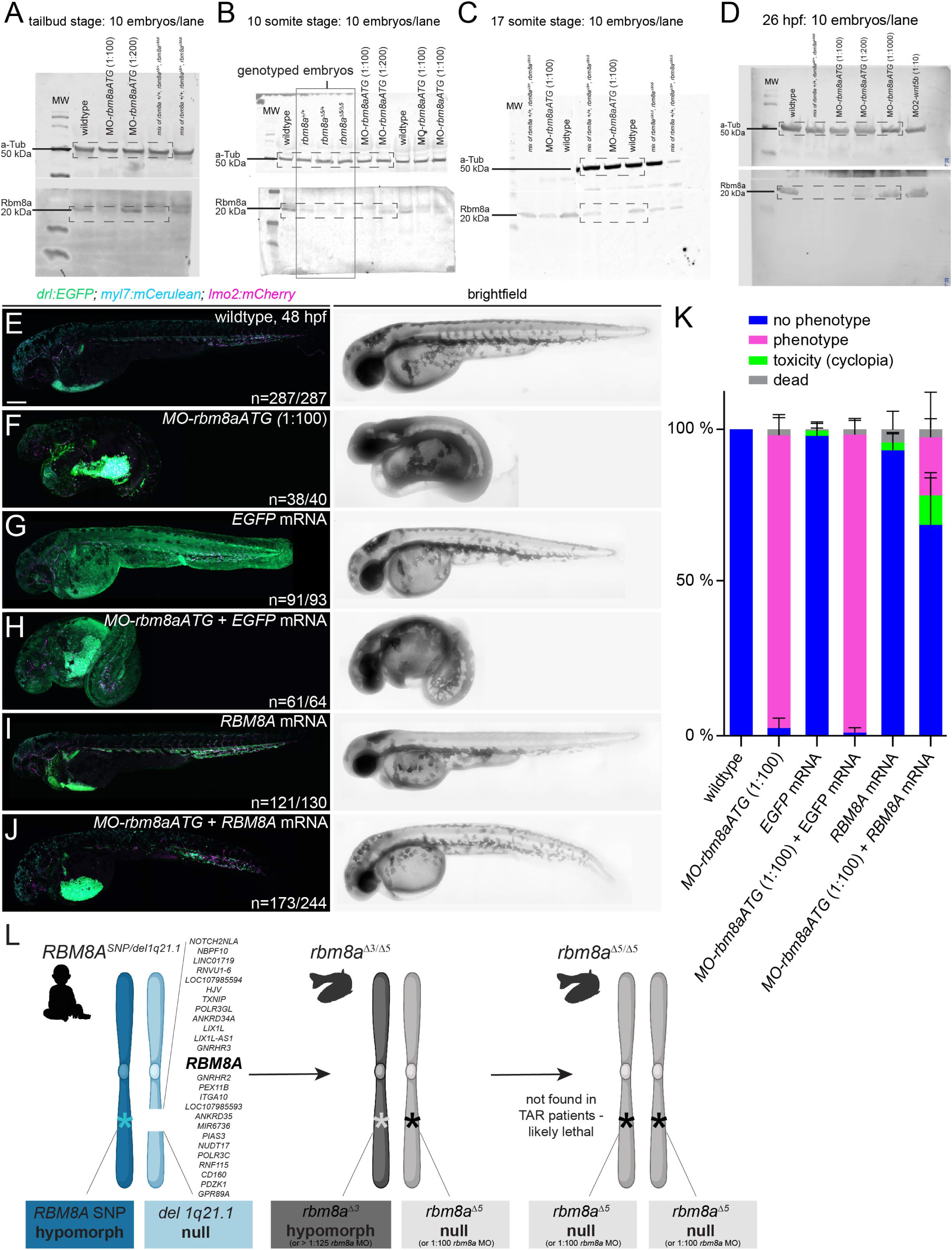
Human *RBM8A* mRNA rescues the *rbm8a* morpholino phenotype. (**A-D**) Uncropped images of western blots displayed in Fig. 1B. Dashed lines represent frames included in Fig. 1B. MW: molecular weight marker. 50 kDa a-Tubulin: loading control. (**A**) Ten tailbud stage embryos per well. 20 kD a-Rbm8a shows 2 bands at this timepoint, with the lower one presumably being the specific one, the upper one phosphorylated protein. (**B**) Ten 10 somite stage embryos per lane. Box indicates rbm8a mutant embryos genotyped individually prior to pooling ten embryos of respective genotype. (**C**) Lanes 2-4: 30 17 hpf embryos per lane; lanes 5-9: Ten 17 hpf embryos per lane; lane 10: Ten 26 hpf embryos per lane; lanes 4-10 loading control 50 kDa a-Tubulin. (**D**) Ten 26 hpf embryos per lane. (**E-J**) Representative confocal images of *drl:EGFP; myl7:mCerulean;lmo2:mCherry* wildtype, morphant, rescued and control injected zebrafish embryos at 48 hpf with brightfield images for reference on the right. (**E**) Representative image of wildtype embryo. (**F**) Representative image of MO-*rbm8a*ATG injected embryo. (**G**) Representative image of *EGFP* mRNA injected control embryo. (**H**) Representative image of MO-*rbm8a*ATG and *EGFP* mRNA co-injected control embryo. (**I**) Representative image of *RBM8A* mRNA injected control embryo. (**J** Representative image of MO-*rbm8a*ATG and *RBM8A* mRNA co-injected (rescued) embryo. (**K**) Overview of rescue. Note how nearly 100% of MO-*rbm8a*ATG groups and only 25% of the rescued embryos have a phenotype. Datapoints (%) shown with mean and standard deviation (see **Supplementary Data 1** for details). (**L**) Compound heterozygosity for *1q21.1* microdeletions and polymorphisms in human *RBM8A* as observed in TAR syndrome patients, and *rbm8a* mutant allele combinations in zebrafish. Created with Biorender.com (Subscription: Individual; Agreement number: TT25WRFMB0). Scale bar in **E**: 200 μm (applies to panels **E-J** and corresponding brightfield images).

**Supplementary Figure 2:**
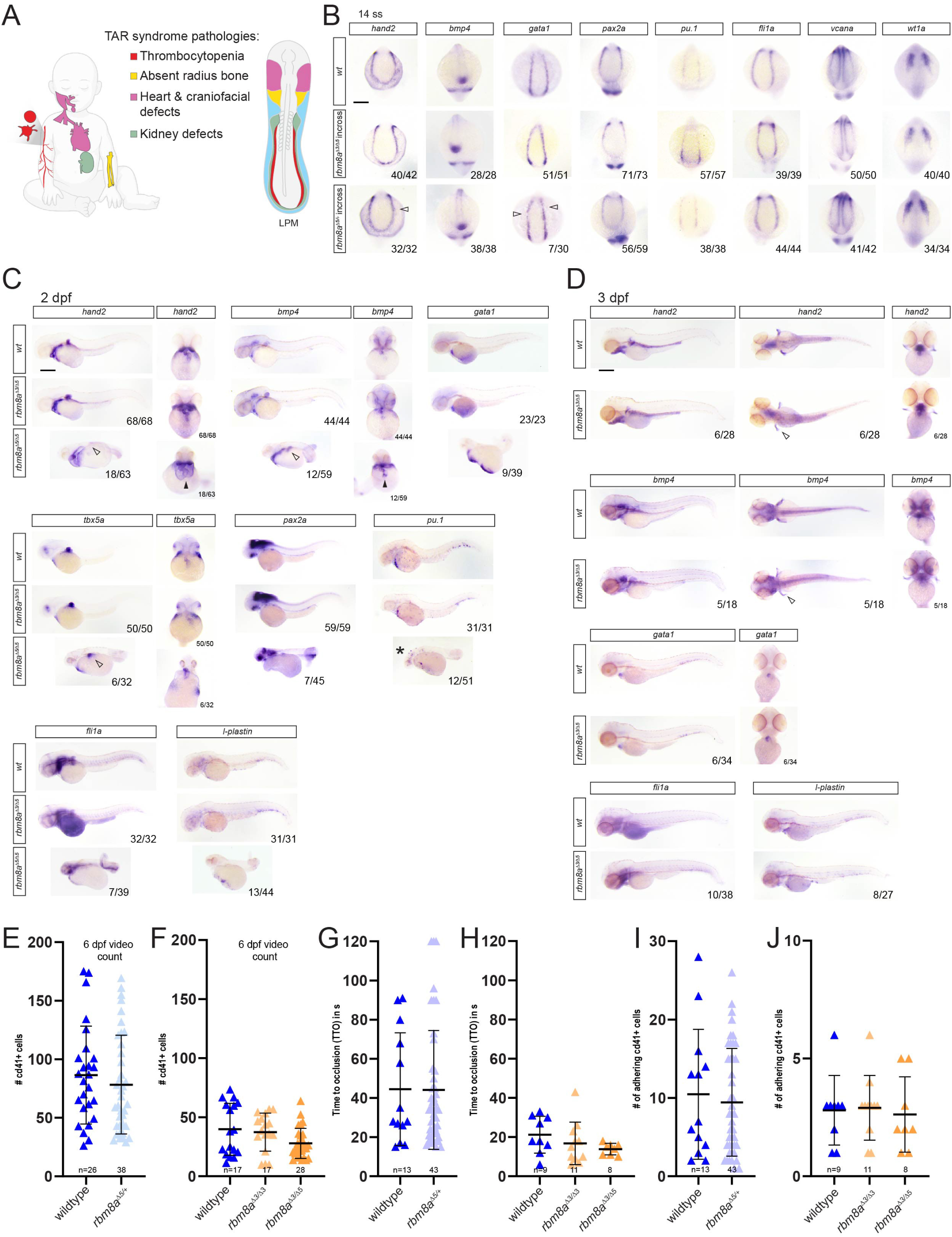
*cd41*:EGFP-expressing thrombocyte quantification using video capture and laser injury assay. (**A**) Hypothesized connection of the human TAR syndrome phenotypes with early lateral plate mesoderm (LPM). Created with Biorender.com (Subscription: Individual; Agreement number: AO25WQRSMG). (**B-D**) *rbm8a*-mutant zebrafish do not show overt LPM defects in early development. mRNA *in situ* hybridization (ISH) to document gene expression of individual LPM-associated genes that demarcate critical cell fates and structures. 3 dpf only shown for *rbm8a^Δ3/Δ5^* hypomorphs as *rbm8a^Δ5/Δ5^* do not survive until that timepoint. Numbers indicate embryos from depicted phenotype within a representative clutch, with confirmed genotyping as control. See text for details. *rbm8a*-mutant embryos show no appreciable gross anomalies in LPM formation, patterning, and individual structures as per these markers, indicating no pleiotropic LPM defects. Individual embryos with mutant *rbm8a* allele combinations showed small disruptions of the initially bilateral LPM stripes with ISH for *gata1* and *hand2* (arrowheads in **B**), but with variable penetrance. While starting functional blood circulation (*gata1*-positive erythrocytes over the yolk, **C**), *pu.1*-positive myeloid cells predominantly infiltrate the brain region with increasing age, consistent with previously reported onset of neuronal apoptosis and necrosis (**C**, asterisk). *rbm8a*-perturbed zebrafish form kidney progenitors from properly patterned *pax2a*-positive bilateral stripes (**B,C**). *rbm8a*-perturbed zebrafish also form pectoral fins and (despite circulation defects in *rbm8a^Δ5/Δ5^*) two-chambered, looped hearts with pericardia akin to wild type references (*tbx5a*, *hand2*, *bmp4*, **C**, and *hand2*, *bmp4*, **D**; black arrowheads: hearts, clear arrowheads: pectoral fins). (**E,F**) Zebrafish with respective genotypes were incrossed and larvae collected. At 6 dpf, the number of circulating *cd41:EGFP*-positive thrombocytes was measured (video count). No statistically significant differences (Mann Whitney test) were detected between the video count method and the semi-automated count method as documented in Fig. 2 (**E**). The number of circulating *cd41:EGFP*-positive thrombocytes demonstrated a trending yet not significant reduction in 6 dpf *rbm8a^Δ3/Δ3^* and *rbm8a^Δ3/Δ5^* zebrafish larvae, as shown with the semi-automated count method in Fig. 2K (**F**). (**G-J**) Time to occlusion (TTO) after laser-mediated arterial endothelial injury and number of *cd41:EGFP*-labeled thrombocytes adhering to the site of occlusion. 6 dpf larvae were subjected to laser-mediated arterial endothelial injury, time to occlusion (TTO) was measured up to 120 seconds, and the number of *cd41:EGFP*-labeled thrombocytes adherent to the site of occlusion were counted over 120 seconds, followed by genotyping. The numbers above the x-axis indicate the number of larvae tested. There were no statistically significant differences in TTO of wildtype compared to heterozygous *rbm8a^Δ5/+^* (**G**), homozygous *rbm8a^Δ3/Δ3^* or trans-heterozygous *rbm8a^Δ3/Δ5^* zebrafish larvae (**H**), or the number of adhering thrombocytes (**I,J**). Significance calculated by Mann-Whitney test (see **Supplementary Data 2** for details). Scale bar in **B:** 250 μm (applies to all panels in **B**), scale bars in **C,D**: 200 μm (applies to all panels in **C-D**).

**Supplementary Figure 3:**
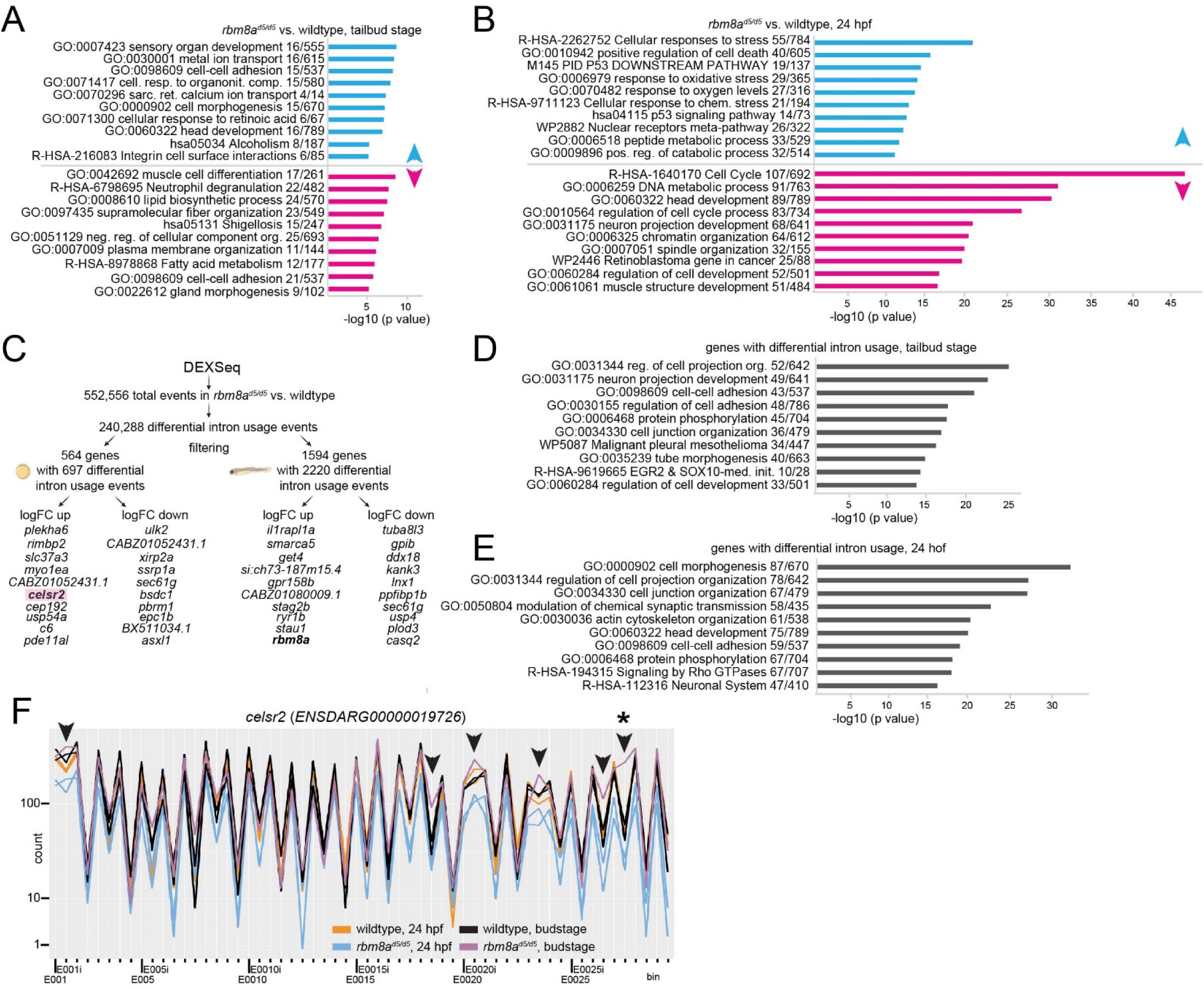
Quantitative and qualitative transcriptome analysis of zebrafish *rbm8a* mutants. (**A,B**) Disrupted (up- and down-regulated) pathways and processes in *rbm8a*-mutant zebrafish embryos at tailbud stage (**A**) and at 24 hpf (**B**); blue indicates upregulation, magenta indicates downregulation (corresponding arrowheads genes (see **Supplementary Data 5** for details). (**C**) DEXSeq analysis to identify genes with differential intron usage reveals transcripts with defects that are retained in rbm8a-mutant embryos. Workflow of the analysis, output, and top altered genes (see **Supplementary Data 6 and 7** for details) (**D,E**) Disrupted (up- and down-regulated) pathways and processes of all genes with intron retention from (**C**) at tailbud stage (**D**) and at 24 hpf (**E**) (see **Supplementary Data 8** for details). (**F**) Read count plot for *celsr2* as representative gene that shows retained intron events. Arrowheads indicate increased intron reads, asterisk indicates majority of transcript with retained individual intron.

## Notes

### Summary of Updates

The newly revised manuscript includes several text edits for clarity and a revised title to better reflect the work's focus. The revisions also incorporate a thorough re-count of all quantitative cell measurements, additional in situ panels to document gene expression, and expanded supplementary data.

## REFERENCES

1. Petit, F. & Boussion, S. Thrombocytopenia Absent Radius Syndrome. GeneReviews® (2022).

2. Albers, C. A., Newbury-Ecob, R., Ouwehand, W. H. & Ghevaert, C. New insights into the genetic basis of TAR (thrombocytopenia-absent radii) syndrome. Curr Opin Genet Dev 23, 316–23 (2013).

3. Klopocki, E. et al. Complex inheritance pattern resembling autosomal recessive inheritance involving a microdeletion in thrombocytopenia-absent radius syndrome. Am J Hum Genet 80, 232–240 (2007).

4. Greenhalgh, K. L. Thrombocytopenia-absent radius syndrome: a clinical genetic study. J Med Genet 39, 876–881 (2002).

5. Boussion, S. et al. TAR syndrome: clinical and molecular characterization of a cohort of 26 patients and description of novel non-coding variants of RBM8A. Hum Mutat 41, 1220–1225 (2020).

6. Albers, C. A. et al. Compound inheritance of a low-frequency regulatory SNP and a rare null mutation in exon-junction complex subunit RBM8A causes TAR syndrome. Nat Genet 44, 435–9, S1-2 (2012).

7. Bottillo, I. et al. Prenatal diagnosis and post-mortem examination in a fetus with thrombocytopenia-absent radius (TAR) syndrome due to compound heterozygosity for a 1q21.1 microdeletion and a RBM8A hypomorphic allele: a case report. BMC Res Notes 6, 376 (2013).

8. Zhao, X. F., Nowak, N. J., Shows, T. B. & Aplan, P. D. MAGOH Interacts with a Novel RNA-Binding Protein. Genomics 63, 145–148 (2000).

9. Chuang, T.-W., Lee, K.-M. & Tarn, W.-Y. Function and pathological implications of exon junction complex factor Y14. Biomolecules 5, 343–55 (2015).

10. Ashton-Beaucage, D. et al. The Exon Junction Complex Controls the Splicing of mapk and Other Long Intron-Containing Transcripts in Drosophila. Cell 143, 251–262 (2010).

11. Roignant, J. Y. & Treisman, J. E. Exon Junction Complex Subunits Are Required to Splice Drosophila MAP Kinase, a Large Heterochromatic Gene. Cell 143, 238–250 (2010).

12. Hir, H. Le, Saulière, J. & Wang, Z. The exon junction complex as a node of post-transcriptional networks. Nature Reviews Molecular Cell Biology 2015 17:1 17, 41–54 (2015).

13. Kurosaki, T., Popp, M. W. & Maquat, L. E. Quality and quantity control of gene expression by nonsense-mediated mRNA decay. Nature Reviews Molecular Cell Biology 2019 20:7 20, 406–420 (2019).

14. Gangras, P. et al. Zebrafish rbm8a and magoh mutants reveal EJC developmental functions and new 3′UTR intron-containing NMD targets. PLoS Genet 16, e1008830 (2020).

15. Schier, A. F. & Talbot, W. S. Molecular genetics of axis formation in zebrafish. Annu Rev Genet 39, 561–613 (2005).

16. Onimaru, K., Shoguchi, E., Kuratani, S. & Tanaka, M. Development and evolution of the lateral plate mesoderm: comparative analysis of amphioxus and lamprey with implications for the acquisition of paired fins. Dev Biol 359, 124–136 (2011).

17. Schoenwolf, G. C., Garcia-Martinez, V. & Dias, M. S. Mesoderm movement and fate during avian gastrulation and neurulation. Dev Dyn 193, 235–48 (1992).

18. Garcia-Martinez, V. & Schoenwolf, G. C. Positional control of mesoderm movement and fate during avian gastrulation and neurulation. Dev Dyn 193, 249–56 (1992).

19. Davidson, A. J. & Zon, L. I. The ‘definitive’ (and ’primitive’) guide to zebrafish hematopoiesis. Oncogene 23, 7233–7246 (2004).

20. Prummel, K. D., Nieuwenhuize, S. & Mosimann, C. The lateral plate mesoderm. 147, (2020).

21. Kocere, A., Lalonde, R. L., Mosimann, C. & Burger, A. Lateral thinking in syndromic congenital cardiovascular disease. Dis Model Mech 16, (2023).

22. Olson, O. C., Kang, Y. A. & Passegué, E. Normal Hematopoiesis Is a Balancing Act of Self-Renewal and Regeneration. Cold Spring Harb Perspect Med 10, 1–23 (2020).

23. Rieger, M. A. & Schroeder, T. Hematopoiesis. Cold Spring Harb Perspect Biol 4, (2012).

24. de Pater, E. & Trompouki, E. Bloody zebrafish: Novel methods in normal and malignant hematopoiesis. Front Cell Dev Biol (2018) doi:10.3389/fcell.2018.00124.

25. Orkin, S. H. & Zon, L. I. Hematopoiesis: an evolving paradigm for stem cell biology. Cell 132, 631–644 (2008).

26. Bianchi, E., Norfo, R., Pennucci, V., Zini, R. & Manfredini, R. Genomic landscape of megakaryopoiesis and platelet function defects. Blood Preprint at 10.1182/blood-2015-07-607952 (2016).

27. Machlus, K. R. & Italiano, J. E. The incredible journey: From megakaryocyte development to platelet formation. Journal of Cell Biology Preprint at 10.1083/jcb.201304054 (2013).

28. Tober, J. et al. The megakaryocyte lineage originates from hemangioblast precursors and is an integral component both of primitive and of definitive hematopoiesis. Blood 109, 1433 (2007).

29. Trowbridge, J. J., Xenocostas, A., Moon, R. T. & Bhatia, M. Glycogen synthase kinase-3 is an in vivo regulator of hematopoietic stem cell repopulation. Nat Med 12, 89–98 (2006).

30. Yzaguirre, A. D., de Bruijn, M. F. T. R. & Speck, N. A. The role of Runx1 in embryonic blood cell formation. in Advances in Experimental Medicine and Biology (2017). doi:10.1007/978-981-10-3233-2_4.

31. Kissa, K. & Herbomel, P. Blood stem cells emerge from aortic endothelium by a novel type of cell transition. Nature 464, 112–115 (2010).

32. Gao, L. et al. RUNX1 and the endothelial origin of blood. Exp Hematol (2018) doi:10.1016/j.exphem.2018.10.009.

33. Chen, M. J., Yokomizo, T., Zeigler, B. M., Dzierzak, E. & Speck, N. A. Runx1 is required for the endothelial to haematopoietic cell transition but not thereafter. Nature 457, 887–891 (2009).

34. Shooshtarizadeh, P. et al. Gfi1b regulates the level of Wnt/β-catenin signaling in hematopoietic stem cells and megakaryocytes. Nat Commun 10, 1270 (2019).

35. Frame, J. M., McGrath, K. E. & Palis, J. Erythro-Myeloid Progenitors: “definitive” hematopoiesis in the conceptus prior to the emergence of hematopoietic stem cells. Blood Cells Mol Dis 51, 220–225 (2013).

36. Palis, J., Robertson, S., Kennedy, M., Wall, C. & Keller, G. Development of erythroid and myeloid progenitors in the yolk sac and embryo proper of the mouse. Development 126, 5073–84 (1999).

37. Beauchemin, H. & Möröy, T. Multifaceted Actions of GFI1 and GFI1B in Hematopoietic Stem Cell Self-Renewal and Lineage Commitment. Front Genet 11, 1268 (2020).

38. Cheng, A. N., Bao, E. L., Fiorini, C. & Sankaran, V. G. Macrothrombocytopenia associated with a rare GFI1B missense variant confounding the presentation of immune thrombocytopenia. Pediatr Blood Cancer 66, e27874 (2019).

39. Saleque, S., Cameron, S. & Orkin, S. H. The zinc-finger proto-oncogene Gfi-1b is essential for development of the erythroid and megakaryocytic lineages. Genes Dev 16, 301–6 (2002).

40. Cooney, J. D. et al. Teleost growth factor independence (gfi) genes differentially regulate successive waves of hematopoiesis. Dev Biol 373, 431–441 (2013).

41. Moore, C. et al. Gfi1aa and Gfi1b set the pace for primitive erythroblast differentiation from hemangioblasts in the zebrafish embryo. Blood Adv 2, 2589–2606 (2018).

42. Thambyrajah, R. et al. A gene trap transposon eliminates haematopoietic expression of zebrafish Gfi1aa, but does not interfere with haematopoiesis. Dev Biol 417, 25–39 (2016).

43. Shin, M., Nagai, H. & Sheng, G. Notch mediates Wnt and BMP signals in the early separation of smooth muscle progenitors and blood/endothelial common progenitors. Development 136, 595–603 (2009).

44. Nostro, M. C., Cheng, X., Keller, G. M. & Gadue, P. Wnt, Activin, and BMP Signaling Regulate Distinct Stages in the Developmental Pathway from Embryonic Stem Cells to Blood. Cell Stem Cell 2, 60–71 (2008).

45. Woll, P. S. et al. Wnt signaling promotes hematoendothelial cell development from human embryonic stem cells. Blood 111, 122–131 (2008).

46. Tran, H. T., Sekkali, B., Van Imschoot, G., Janssens, S. & Vleminckx, K. Wnt/beta-catenin signaling is involved in the induction and maintenance of primitive hematopoiesis in the vertebrate embryo. Proc Natl Acad Sci U S A 107, 16160–5 (2010).

47. Kelly, G. M., Greenstein, P., Erezyilmaz, D. F. & Moon, R. T. Zebrafish wnt8 and wnt8b share a common activity but are involved in distinct developmental pathways. Development 121, 1787–99 (1995).

48. Jones, C. & Chen, P. Planar cell polarity signaling in vertebrates. BioEssays 29, 120–132 (2007).

49. Butler, M. T. & Wallingford, J. B. Planar cell polarity in development and disease. Nature Reviews Molecular Cell Biology 2017 18:6 18, 375–388 (2017).

50. Mlodzik, M. Planar cell polarization: do the same mechanisms regulate Drosophila tissue polarity and vertebrate gastrulation? Trends in Genetics 18, 564–571 (2002).

51. Williams, M. L. K. & Solnica-Krezel, L. Nodal and planar cell polarity signaling cooperate to regulate zebrafish convergence & extension gastrulation movements. Elife (2020) doi:10.7554/eLife.54445.

52. Merks, A. M. et al. Planar cell polarity signalling coordinates heart tube remodelling through tissue-scale polarisation of actomyosin activity. Nat Commun 9, 2161 (2018).

53. Creighton, J. H. & Jessen, J. R. Core pathway proteins and the molecular basis of planar polarity in the zebrafish gastrula. Semin Cell Dev Biol (2021) doi:10.1016/J.SEMCDB.2021.09.015.

54. Li, D., Angermeier, A. & Wang, J. Planar cell polarity signaling regulates polarized second heart field morphogenesis to promote both arterial and venous pole septation. Development (Cambridge) 146, (2019).

55. Roszko, I., Sawada, A. & Solnica-Krezel, L. Regulation of convergence and extension movements during vertebrate gastrulation by the Wnt/PCP pathway. Semin Cell Dev Biol 20, 986–97 (2009).

56. Williams, M. L. K. et al. Gon4l regulates notochord boundary formation and cell polarity underlying axis extension by repressing adhesion genes. Nat Commun 9, 1319 (2018).

57. Humphries, A. C., Narang, S. & Mlodzik, M. Mutations associated with human neural tube defects display disrupted planar cell polarity in drosophila. Elife 9, (2020).

58. Goggolidou, P. et al. Atmin mediates kidney morphogenesis by modulating Wnt signaling. Hum Mol Genet 23, 5303–5316 (2014).

59. Chen, Z. et al. Genetic analysis of Wnt/PCP genes in neural tube defects. BMC Med Genomics 11, (2018).

60. Jessen, J. R. et al. Zebrafish trilobite identifies new roles for Strabismus in gastrulation and neuronal movements. Nat Cell Biol 4, 610–615 (2002).

61. Marlow, F., Topczewski, J., Sepich, D. & Solnica-Krezel, L. Zebrafish Rho kinase 2 acts downstream of Wnt11 to mediate cell polarity and effective convergence and extension movements. Curr Biol 12, 876– 884 (2002).

62. Clements, W. K. et al. A somitic Wnt16/Notch pathway specifies haematopoietic stem cells. Nature 474, 220–225 (2011).

63. Genthe, J. R. & Clements, W. K. R-spondin 1 is required for specification of hematopoietic stem cells through Wnt16 and Vegfa signaling pathways. Development 144, 590 (2017).

64. Sugimura, R. et al. Noncanonical Wnt signaling maintains hematopoietic stem cells in the niche. Cell 150, 351–365 (2012).

65. Bono, F., Ebert, J., Lorentzen, E. & Conti, E. The Crystal Structure of the Exon Junction Complex Reveals How It Maintains a Stable Grip on mRNA. Cell 126, 713–725 (2006).

66. Fribourg, S., Gatfield, D., Izaurralde, E. & Conti, E. A novel mode of RBD-protein recognition in the Y14– Mago complex. Nature Structural & Molecular Biology 2003 10:6 10, 433–439 (2003).

67. Shi, H. & Xu, R. M. Crystal structure of the Drosophila Mago nashi–Y14 complex. Genes Dev 17, 971– 976 (2003).

68. Stainier, D. Y. R. et al. Guidelines for morpholino use in zebrafish. PLoS Genet 13, e1007000 (2017).

69. Lin, H. F. et al. Analysis of thrombocyte development in CD41-GFP transgenic zebrafish. Blood (2005) doi:10.1182/blood-2005-01-0179.

70. Belmonte, R. L. et al. son is necessary for proper vertebrate blood development. PLoS One 16, e0247489 (2021).

71. Gansner, J. M. et al. Sorting zebrafish thrombocyte lineage cells with a Cd41 monoclonal antibody enriches hematopoietic stem cell activity. Blood blood-2016-12-759993 (2017) doi:10.1182/blood-2016-12-759993.

72. Bertrand, J. Y., Kim, A. D., Teng, S. & Traver, D. CD41+ cmyb+ precursors colonize the zebrafish pronephros by a novel migration route to initiate adult hematopoiesis. Development 135, 1853–1862 (2008).

73. Huarng, M. C. & Shavit, J. A. Simple and Rapid Quantification of Thrombocytes in Zebrafish Larvae. Zebrafish 12, 238 (2015).

74. Rost, M. S., Grzegorski, S. J. & Shavit, J. A. Quantitative methods for studying hemostasis in zebrafish larvae. Methods Cell Biol 134, 377–389 (2016).

75. Jagadeeswaran, P., Carrillo, M., Radhakrishnan, U. P., Rajpurohit, S. K. & Kim, S. Laser-induced thrombosis in zebrafish. Methods Cell Biol 101, 197–203 (2011).

76. Zhou, Y. et al. Metascape provides a biologist-oriented resource for the analysis of systems-level datasets. Nat Commun 10, (2019).

77. McMahon, J. J., Miller, E. E. & Silver, D. L. The exon junction complex in neural development and neurodevelopmental disease. International Journal of Developmental Neuroscience Preprint at 10.1016/j.ijdevneu.2016.03.006 (2016).

78. Palacios, I. M., Gatfield, D., St Johnston, D. & Izaurralde, E. An eIF4AIII-containing complex required for mRNA localization and nonsense-mediated mRNA decay. Nature 427, 753–7 (2004).

79. Anders, S., Reyes, A. & Huber, W. Detecting differential usage of exons from RNA-seq data. Genome Res 22, 2008–2017 (2012).

80. Joshi, B., Gaur, H., Hui, S. P. & Patra, C. Celsr family genes are dynamically expressed in embryonic and juvenile zebrafish. Dev Neurobiol 82, 192–213 (2022).

81. Dooley, K. a, Davidson, A. J. & Zon, L. I. Zebrafish scl functions independently in hematopoietic and endothelial development. Dev Biol 277, 522–36 (2005).

82. Zhang, X. Y. & Rodaway, A. R. F. SCL-GFP transgenic zebrafish: In vivo imaging of blood and endothelial development and identification of the initial site of definitive hematopoiesis. Dev Biol 307, 179–194 (2007).

83. Love, A. M., Prince, D. J. & Jessen, J. R. Vangl2-dependent regulation of membrane protrusions and directed migration requires a fibronectin extracellular matrix. Development dev.165472 (2018) doi:10.1242/dev.165472.

84. Roszko, I., Sepich, D. S., Jessen, J. R., Chandrasekhar, A. & Solnica-Krezel, L. A dynamic intracellular distribution of Vangl2 accompanies cell polarization during zebrafish gastrulation. Development (Cambridge) (2015) doi:10.1242/dev.119032.

85. Yang, Y. & Mlodzik, M. Wnt-Frizzled/Planar Cell Polarity Signaling: Cellular Orientation by Facing the Wind (Wnt). Annu Rev Cell Dev Biol 31, 623–646 (2015).

86. Prince, D. J. & Jessen, J. R. Dorsal convergence of gastrula cells requires a Vangl2 and adhesion protein-dependent change in protrusive activity. Development (2019) doi:10.1242/dev.182188.

87. Marlow, F. et al. Functional interactions of genes mediating convergent extension, knypek and trilobite, during the partitioning of the eye primordium in zebrafish. Dev Biol 203, 382–399 (1998).

88. Li, X. et al. Gpr125 modulates Dishevelled distribution and planar cell polarity signaling. Development (Cambridge) 140, 3028–3039 (2013).

89. Hammerschmidt, M. et al. Mutations affecting morphogenesis during gastrulation and tail formation in the zebrafish, Danio rerio. Development 123, 143–51 (1996).

90. Solnica-Krezel, L. et al. Mutations affecting cell fates and cellular rearrangements during gastrulation in zebrafish. Development 123, 67–80 (1996).

91. Whitfield, T. T. et al. Mutations affecting development of the zebrafish inner ear and lateral line. Development 123, 241–54 (1996).

92. Jussila, M., Boswell, C. W., Griffiths, N. W., Pumputis, P. G. & Ciruna, B. Live imaging and conditional disruption of native PCP activity using endogenously tagged zebrafish sfGFP-Vangl2. Nature Communications 2022 13:1 13, 1–13 (2022).

93. Heisenberg, C. P. et al. Silberblick/Wnt11 mediates convergent extension movements during zebrafish gastrulation. Nature 405, 76–81 (2000).

94. Pierpont, M. E. et al. Genetic Basis for Congenital Heart Disease: Revisited: A Scientific Statement From the American Heart Association. Circulation 138, (2018).

95. Kilian, B. et al. The role of Ppt/Wnt5 in regulating cell shape and movement during zebrafish gastrulation. Mech Dev 120, 467–476 (2003).

96. Topczewski, J. et al. The zebrafish glypican knypek controls cell polarity during gastrulation movements of convergent extension. Dev Cell 1, 251–264 (2001).

97. Schulte-Merker, S. et al. Expression of zebrafish goosecoid and no tail gene products in wild-type and mutant no tail embryos. Development 120, 843–852 (1994).

98. Schulte-Merker, S., van Eeden, F. J., Halpern, M. E., Kimmel, C. B. & Nüsslein-Volhard, C. no tail (ntl) is the zebrafish homologue of the mouse T (Brachyury) gene. Development 120, 1009–15 (1994).

99. Weinberg, E. S. et al. Developmental regulation of zebrafish MyoD in wild-type, no tail and spadetail embryos. 122, 271–280 (1996).

100. Akimenko, M. A., Ekker, M., Wegner, J., Lin, W. & Westerfield, M. Combinatorial expression of three zebrafish genes related to distal-less: part of a homeobox gene code for the head. J Neurosci 14, 3475– 3486 (1994).

101. Vogel, A. M. & Gerster, T. Expression of a zebrafish Cathepsin L gene in anterior mesendoderm and hatching gland. Dev Genes Evol 477–479 (1997) doi:10.1007/S004270050078.

102. Mosimann, C. et al. Chamber identity programs drive early functional partitioning of the heart. Nat Commun 6, (2015).

103. Panakova, D., Werdich, A. A. & Macrae, C. A. Wnt11 patterns a myocardial electrical gradient through regulation of the L-type Ca(2+) channel. Nature 466, 874–878 (2010).

104. Herpers, R., Van De Kamp, E., Duckers, H. J. & Schulte-Merker, S. Redundant roles for sox7 and sox18 in arteriovenous specification in Zebrafish. Circ Res 102, 12–15 (2008).

105. Charney, R. M. et al. Foxh1 Occupies cis -Regulatory Modules Prior to Dynamic Transcription Factor Interactions Controlling the Mesendoderm Gene Program. Dev Cell 40, 595–607.e4 (2017).

106. Long, Q. et al. GATA-1 expression pattern can be recapitulated in living transgenic zebrafish using GFP reporter gene. Development 124, 4105–4111 (1997).

107. Wei, W. et al. Gfi1.1 regulates hematopoietic lineage differentiation during zebrafish embryogenesis. Cell Res 18, 677–685 (2008).

108. Kalev-Zylinska, M. L. et al. Runx1 is required for zebrafish blood and vessel development and expression of a human RUNX1-CBF2T1 transgene advances a model for studies of leukemogenesis. Development 129, 2015–2030 (2002).

109. Horsfield, J. A. et al. Cohesin-dependent regulation of Runx genes. Development 134, 2639–2649 (2007).

110. Landowski, M. et al. Novel deletion of RPL15 identified by array-comparative genomic hybridization in Diamond-Blackfan anemia. Hum Genet 132, 1265–1274 (2013).

111. Gazda, H. T. et al. Ribosomal Protein S24 Gene Is Mutated in Diamond-Blackfan Anemia. The American Journal of Human Genetics 79, 1110–1118 (2006).

112. Stoll, C., Dott, B., Alembik, Y. & Roth, M. P. Associated malformations among infants with radial ray deficiency. Genetic Counseling (2013).

113. Thompson, A. A., Woodruff, K., Feig, S. A., Nguyen, L. T. & Carolyn Schanen, N. Congenital thrombocytopenia and radio-ulnar synostosis: a new familial syndrome. Br J Haematol 113, 866–870 (2001).

114. Parmar, K., D’Andrea, A. & Niedernhofer, L. J. Mouse models of Fanconi anemia. Mutat Res 668, 133 (2009).

115. Bhandari, J., Thada, P. K. & Puckett, Y. Fanconi Anemia. StatPearls (2022).

116. Balduini, C. L. The name counts: the case of ‘congenital amegakaryocytic thrombocytopenia’. Haematologica (2022) doi:10.3324/HAEMATOL.2022.282024.

117. Thompson, A. A. & Nguyen, L. T. Amegakaryocytic thrombocytopenia and radio-ulnar synostosis are associated with HOXA11 mutation. Nature Genetics 2000 26:4 26, 397–398 (2000).

118. Niihori, T. et al. Mutations in MECOM, Encoding Oncoprotein EVI1, Cause Radioulnar Synostosis with Amegakaryocytic Thrombocytopenia. 97, 848–854 (2015).

119. Perkins, A. S., Mercer, J. A., Jenkins, N. A. & Copeland, N. G. Patterns of Evi-1 expression in embryonic and adult tissues suggest that Evi-1 plays an important regulatory role in mouse development. Development 111, 479–487 (1991).

120. Davis, A. P., Witte, D. P., Hsieh-Li, H. M., Potter, S. S. & Capecchi, M. R. Absence of radius and ulna in mice lacking hoxa-11 andhoxd-11. Nature 1995 375:6534 375, 791–795 (1995).

121. Patterson, L. T., Pembaur, M. & Potter, S. S. Hoxa11 and Hoxd11 regulate branching morphogenesis of the ureteric bud in the developing kidney. Development 128, 2153–2161 (2001).

122. Yuan, X., Wang, X., Bi, K. & Jiang, G. The role of EVI-1 in normal hematopoiesis and myeloid malignancies (review). Int J Oncol 47, 2028–2036 (2015).

123. Bard-Chapeau, E. A. et al. Ecotopic viral integration site 1 (EVI1) regulates multiple cellular processes important for cancer and is a synergistic partner for FOS protein in invasive tumors. Proc Natl Acad Sci U S A 109, 2168–2173 (2012).

124. Li, J. et al. Limb development genes underlie variation in human fingerprint patterns. Cell 185, 95–112.e18 (2022).

125. Yokomizo, T. et al. Independent origins of fetal liver haematopoietic stem and progenitor cells. Nature 2022 1–6 (2022) doi:10.1038/s41586-022-05203-0.

126. Konantz, M. et al. Evi1 regulates Notch activation to induce zebrafish hematopoietic stem cell emergence. EMBO J 459, 1131–1135 (2016).

127. Mugford, J. W., Sipilä, P., Kobayashi, A., Behringer, R. R. & McMahon, A. P. Hoxd11 specifies a program of metanephric kidney development within the intermediate mesoderm of the mouse embryo. Dev Biol 319, 396–405 (2008).

128. Horvat-Switzer, R. D. & Thompson, A. A. HOXA11 mutation in amegakaryocytic thrombocytopenia with radio-ulnar synostosis syndrome inhibits megakaryocytic differentiation in vitro. Blood Cells Mol Dis 37, 55–63 (2006).

129. Le Hir, H. & Andersen, G. R. Structural insights into the exon junction complex. Curr Opin Struct Biol 18, 112–119 (2008).

130. Pang, H. et al. Disorders Associated With Diverse, Recurrent Deletions and Duplications at 1q21.1. *Front Genet* (2020) doi:10.3389/fgene.2020.00577.

131. Ballmaier, M. et al. Defective c-Mpl signaling in the syndrome of thrombocytopenia with absent radii. Stem Cells 16 **Suppl 2**, 177–184 (1998).

132. Yamanaka, Y., Tamplin, O. J., Beckers, A., Gossler, A. & Rossant, J. Live imaging and genetic analysis of mouse notochord formation reveals regional morphogenetic mechanisms. Dev Cell 13, 884–96 (2007).

133. Warga, R. M. & Kane, D. a. A role for N-cadherin in mesodermal morphogenesis during gastrulation. Dev Biol 310, 211–25 (2007).

134. Kishimoto, N., Cao, Y., Park, A. & Sun, Z. Cystic Kidney Gene seahorse Regulates Cilia-Mediated Processes and Wnt Pathways. Dev Cell 14, 954–961 (2008).

135. Carreira-Barbosa, F. et al. Prickle 1 regulates cell movements during gastrulation and neuronal migration in zebrafish. Development 130, 4037–4046 (2003).

136. Harrington, M. J., Hong, E., Fasanmi, O. & Brewster, R. Cadherin-mediated adhesion regulates posterior body formation. BMC Dev Biol 7, 1–19 (2007).

137. Yin, C. & Solnica-Krezel, L. Convergence and extension movements mediate the specification and fate maintenance of zebrafish slow muscle precursors. Dev Biol 304, 141–55 (2007).

138. Xing, Y. Y. et al. Mutational analysis of dishevelled genes in zebrafish reveals distinct functions in embryonic patterning and gastrulation cell movements. PLoS Genet 14, (2018).

139. Sahai-Hernandez, P. et al. Dermomyotome-derived endothelial cells migrate to the dorsal aorta to support hematopoietic stem cell emergence. Elife 12, (2023).

140. Kobayashi, I. et al. Jam1a–Jam2a interactions regulate haematopoietic stem cell fate through Notch signalling. Nature 2014 512:7514 512, 319–323 (2014).

141. Heazlewood, S. Y. et al. The RNA-binding protein SRSF3 has an essential role in megakaryocyte maturation and platelet production. Blood 139, 1359–1373 (2022).

142. Bonsi, L. et al. Thrombocytopenia with absent radii (TAR) syndrome: from hemopoietic progenitor to mesenchymal stromal cell disease? Exp Hematol 37, 1–7 (2009).

143. Zhang, Y., Gao, S., Xia, J. & Liu, F. Hematopoietic Hierarchy – An Updated Roadmap. Trends in Cell Biology Preprint at 10.1016/j.tcb.2018.06.001 (2018).

144. Pinho, S. & Frenette, P. S. Haematopoietic Stem Cell Activity and Interactions with the Niche. Nature Reviews Molecular Cell Biology vol. 20 303–320 (Nature Publishing Group, 2019).

145. Doulatov, S., Notta, F., Laurenti, E. & Dick, J. E. Hematopoiesis: A Human Perspective. Cell Stem Cell 10, 120–136 (2012).

146. Fulton, T. et al. Axis Specification in Zebrafish Is Robust to Cell Mixing and Reveals a Regulation of Pattern Formation by Morphogenesis. Current Biology 30, 2984–2994.e3 (2020).

147. Li, D. et al. Spatial regulation of cell cohesion by Wnt5a during second heart field progenitor deployment. Dev Biol (2016) doi:10.1016/j.ydbio.2016.02.017.

148. Gao, B. & Yang, Y. Planar Cell Polarity in vertebrate limb morphogenesis. Curr Opin Genet Dev 23, 438 (2013).

149. Wang, B., Sinha, T., Jiao, K., Serra, R. & Wang, J. Disruption of PCP signaling causes limb morphogenesis and skeletal defects and may underlie Robinow syndrome and brachydactyly type B. Hum Mol Genet 20, 271–85 (2011).

150. Li, D. & Wang, J. Planar cell polarity signaling in mammalian cardiac morphogenesis. Pediatr Cardiol 39, 1052 (2018).

151. Aleström, P. et al. Zebrafish: Housing and husbandry recommendations. Lab Anim 54, 213–224 (2020).

152. Westerfield, M. The Zebrafish Book: A Guide for the Laboratory Use of Zebrafish (Danio Rerio). (University of Oregon Press, Eugene, 2007).

153. Chiavacci, E., Kirchgeorg, L., Felker, A., Burger, A. & Mosimann, C. Early frameshift alleles of zebrafish tbx5a that fail to develop the heartstrings phenotype. Matters (Zur) 2017030000, 103168 (2017).

154. Wang, L. et al. Functional characterization of Lmo2-Cre transgenic zebrafish. Dev Dyn 237, 2139–2146 (2008).

155. Zhou, Y. et al. Latent TGF-βbinding protein 3 identifies a second heart field in zebrafish. Nature 474, (2011).

156. Bassett, A. R., Tibbit, C., Ponting, C. P. P. & Liu, J. L. J.-L. Highly efficient targeted mutagenesis of Drosophila with the CRISPR/Cas9 system. Cell Rep 4, 220–8 (2013).

157. Gagnon, J. A. et al. Efficient mutagenesis by Cas9 protein-mediated oligonucleotide insertion and large-scale assessment of single-guide RNAs. PLoS One 9, e98186 (2014).

158. Burger, A. et al. Maximizing mutagenesis with solubilized CRISPR-Cas9 ribonucleoprotein complexes. Development 143, 2025–37 (2016).

159. Montague, T. G., Cruz, J. M., Gagnon, J. A., Church, G. M. & Valen, E. CHOPCHOP: a CRISPR/Cas9 and TALEN web tool for genome editing. Nucleic Acids Res 42, W401–W407 (2014).

160. Turner, D. L. & Weintraub, H. Expression of achaete-scute homolog 3 in Xenopus embryos converts ectodermal cells to a neural fate. Genes Dev 8, 1434–1447 (1994).

161. Kwan, K. M. et al. The Tol2kit: a multisite gateway-based construction kit for Tol2 transposon transgenesis constructs. Dev Dyn 236, 3088–3099 (2007).

162. Kim, H. J. et al. Wnt5 signaling in vertebrate pancreas development. BMC Biol 3, 23 (2005).

163. Takamiya, M. & Campos-Ortega, J. A. Hedgehog signalling controls zebrafish neural keel morphogenesis via its level-dependent effects on neurogenesis. Developmental Dynamics 235, 978– 997 (2006).

164. Matsui, T. et al. Noncanonical Wnt signaling regulates midline convergence of organ primordia during zebrafisn development. Genes Dev (2005) doi:10.1101/gad.1253605.

165. Fong, S. H., Emelyanov, A., Teh, C. & Korzh, V. Wnt signalling mediated by Tbx2b regulates cell migration during formation of the neural plate. Development 132, 3587–3596 (2005).

166. Schindelin, J., et al. Fiji: an open-source platform for biological-image analysis. Nat Methods 9, 676–82 (2012).

167. Ewels, P., Magnusson, M., Lundin, S. & Käller, M. MultiQC: summarize analysis results for multiple tools and samples in a single report. Bioinformatics 32, 3047–3048 (2016).

168. Patro, R., Duggal, G., Love, M. I., Irizarry, R. A. & Kingsford, C. Salmon provides fast and bias-aware quantification of transcript expression. Nature Methods 2017 14:4 14, 417–419 (2017).

169. Aken, B. L. et al. Ensembl 2017. Nucleic Acids Res 45, D635–D642 (2017).

170. Soneson, C., Love, M. I. & Robinson, M. D. Differential analyses for RNA-seq: transcript-level estimates improve gene-level inferences. 4, 1521 (2016).

171. Robinson, M. D., McCarthy, D. J. & Smyth, G. K. edgeR: a Bioconductor package for differential expression analysis of digital gene expression data. Bioinformatics 26, 139–140 (2010).

172. Lun, A. T. L., Chen, Y. & Smyth, G. K. It’s DE-licious: A Recipe for Differential Expression Analyses of RNA-seq Experiments Using Quasi-Likelihood Methods in edgeR. Methods Mol Biol 1418, 391–416 (2016).

173. Benjamini, Y. & Hochberg, Y. Controlling the False Discovery Rate: A Practical and Powerful Approach to Multiple Testing. Journal of the Royal Statistical Society: Series B (Methodological) 57, 289–300 (1995).

174. Dobin, A. et al. STAR: ultrafast universal RNA-seq aligner. Bioinformatics 29, 15–21 (2013).

175. Quinlan, A. R. & Hall, I. M. BEDTools: a flexible suite of utilities for comparing genomic features. Bioinformatics 26, 841–842 (2010).

176. Kent, W. J., Zweig, A. S., Barber, G., Hinrichs, A. S. & Karolchik, D. BigWig and BigBed: enabling browsing of large distributed datasets. Bioinformatics 26, 2204 (2010).

177. Hahne, F. & Ivanek, R. Visualizing genomic data using Gviz and bioconductor. Methods in Molecular Biology 1418, 335–351 (2016).

178. Soneson, C., Matthes, K. L., Nowicka, M., Law, C. W. & Robinson, M. D. Isoform prefiltering improves performance of count-based methods for analysis of differential transcript usage. 17, 12 (2016).

179. Hörl, D. et al. BigStitcher: reconstructing high-resolution image datasets of cleared and expanded samples. Nat Methods 16, 870–874 (2019).

180. Prummel, K. D. et al. Hand2 delineates mesothelium progenitors and is reactivated in mesothelioma. Nature Communications 2022 13:1 13, 1–21 (2022).

181. Thisse, C. & Thisse, B. High-resolution in situ hybridization to whole-mount zebrafish embryos. Nat Protoc 3, 59–69 (2008).

182. Thompson, M. A. et al. TheclocheandspadetailGenes Differentially Affect Hematopoiesis and Vasculogenesis. Dev Biol 197, 248–269 (1998).

183. North, T. E. et al. Prostaglandin E2 regulates vertebrate haematopoietic stem cell homeostasis. Nature 447, 1007–1011 (2007).

184. Schneider, C. A., Rasband, W. S. & Eliceiri, K. W. NIH Image to ImageJ: 25 years of image analysis. Nature Methods vol. 9 671–675 Preprint at 10.1038/nmeth.2089 (2012).

185. Dobrzycki, T., Krecsmarik, M., Bonkhofer, F., Patient, R. & Monteiro, R. An optimised pipeline for parallel image-based quantification of gene expression and genotyping after in situ hybridisation. Biol Open 7, (2018).

